# Engineering a flexible loop in S-adenosyl-*L*-methionine synthetase enables production of SAM nucleobase analogues with selective biochemical and cellular activity

**DOI:** 10.64898/2026.07.10.737877

**Authors:** Dhritiraj Bastav Kalita, Aditya Bhattacharyya, Vruta Gupte, Vikram Venugopal, Anjali Pattathil, Amrita B. Hazra

**Author notes:** Department of Chemistry and Chemical Biology, Cornell University, Ithaca, NY, 14853, Weill Institute for Cell and Molecular Biology, Cornell University, Ithaca, NY, 14853, United States. Department of Chemistry, Institute of Chemical Technology (ICT), Mumbai, 400019, India. The Stowers Institute for Medical Research, Kansas City, Missouri, 64110, United States (Vikram Venugopal). Institute of Medical Microbiology, RWTH University Hospital, Aachen, 52074, Germany. These authors contributed equally (Dhritiraj Bastav Kalita, Aditya Bhattacharyya).

## Abstract

*S*-adenosyl-*L*-methionine (SAM), an essential cofactor in all forms of life, is synthesized by the enzyme methionine adenosyltransferase (MAT) from methionine and ATP. The adenine moiety in SAM appears to have no direct function in catalysis, and some MAT homologs can utilize natural nucleotide triphosphates *in vitro*, producing the corresponding SAM nucleobase analogues. However, the molecular determinants of nucleotide choice of the MAT enzyme and the cellular significance of the nucleobase in SAM are unclear. In this study, using structure- and bioinformatics-guided mutagenesis, we identify a flexible active-site loop as a major determinant of nucleotide specificity in MAT. Loop mutations and loop swaps convert ATP-selective *Escherichia coli* MAT into variants that accept GTP, CTP, and UTP, enabling enzymatic synthesis and purification of S-guanosyl-, S-cytosyl-, and S-uracyl-L-methionine. Further, we show that these analogues partially rescue the growth of an *E. coli* SAM auxotroph under SAM-limited growth conditions. Biochemical assays show that the analogues bind the tested SAM-utilizing enzymes; they serve as substrates for *E. coli* SAM decarboxylase but do not support detectable methyl transfer by *E. coli* DNA adenine methyltransferase. These results establish the flexible loop as a gatekeeper of MAT nucleotide specificity and show that this loop can be engineered to produce SAM analogues which can selectively participate in downstream cellular metabolism.

**Graphical Abstract/ Table of contents only:** 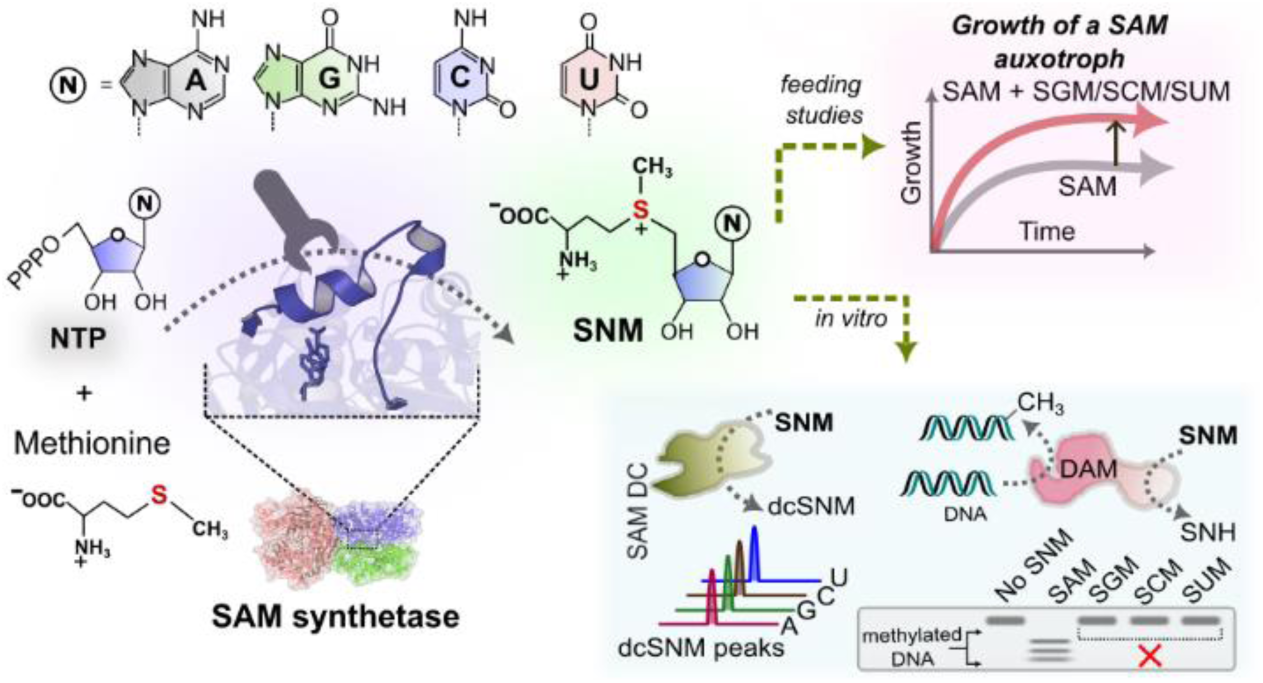

## Introduction

*S*-adenosyl-*L*-methionine (SAM or AdoMet) is an important metabolic cofactor used in a diverse range of biological reactions in primary metabolism across all domains of life.^1,2^ SAM acts broadly as a methyl donor and supports radical-mediated rearrangements.^2,3^ SAM also serves as the precursor of decarboxy-SAM, an intermediate in polyamine biosynthesis. In addition, SAM-dependent enzymes catalyze specialized reactions such as oxygen-independent oxidative decarboxylation in heme biosynthesis, lipid cyclopropanation, radical-based porphyrin ring opening, and radical-promoted self-cyclization.^2,4–6^

Methyl group analogues of SAM have been extensively used as chemical biology probes for site-specific labelling of DNA and proteins.^7–9^ Fluorescent analogues and enzyme inhibitors have also been produced by modifications at the nucleobase and ribose rings.^8^ Despite these efforts, whether replacement of the adenine moiety affects cofactor function in cells remains unclear.^10,11^

SAM synthetase, also known as methionine adenosyltransferase (MAT), catalyzes the bimolecular reaction of ATP and methionine. Several MAT homologs display nucleotide promiscuity and have been used to enzymatically synthesize SAM nucleotide analogues (SNMs), including *S*-guanosyl-*L*-methionine (SGM), *S*-cytosyl-*L*-methionine (SCM), and *S*-uracyl-*L*-methionine (SUM) by using guanosine triphosphate (GTP), cytidine triphosphate (CTP), and uridine triphosphate (UTP), respectively, as well as non-natural analogues such as N^6^-propargyl SAM and N^6^-benzyl SAM.^12–15^ Four MAT homologs have been characterized in literature for their ability to utilize different naturally occurring nucleotides.^12,15–17^ The *Thermococcus kodakarensis* MAT shows activity with ATP, CTP and inosine triphosphate (ITP), and *Cryptosporidium hominis* MAT shows activity with ATP and ITP, forming the corresponding SNMs.^12,16^ The *Methanocaldococcus jannaschii* MAT (*Mj*MAT) and *Homo sapiens* MAT2a (*Hs*MAT2a) can use all four naturally occurring ribonucleotides *in vitro.*^13,15,17,19,20^ In contrast, *Escherichia coli* MAT (*Ec*MAT) is strongly ATP-selective and shows minimal activity with GTP, CTP, or UTP under comparable *in vitro* conditions.^15,18^ The structural basis for this difference in nucleotide specificity remains unclear. Notably, 20 of the 21 active-site residues are identical between *Hs*MAT2a and *Ec*MAT, and making the active sites identical does not alter the nucleotide specificity of either enzyme. This suggests that residues or structural features outside the primary active-site shell contribute to nucleotide substrate choice.^15^

Beyond the enzymatic origin of nucleotide selectivity, a second unresolved question is whether these SAM nucleotide analogues can act as functional cofactors within cells. While SNMs have been tested for their *in vitro* activity with two methyltransferases and two radical SAM enzymes, the functional role of these molecules within cells remains unexplored.^14,21,22^

In this study, we identify a flexible loop near the substrate-binding site as one of the primary determinants of the nucleotide substrate choice for MAT.^20^ Based on a comprehensive structural and evolutionary analysis of MAT, we performed a rational site-directed mutagenesis and loop swap-based engineering strategy, and we converted the ATP-specific *Ec*MAT into variants that are promiscuous for natural nucleotides GTP, CTP and UTP as co-substrates. We further enzymatically synthesized, purified and characterized the SAM analogues SGM, SCM, and SUM, and showed that these can partially rescue the growth of an *E. coli* SAM auxotroph in SAM-limited growth medium. We also showed that these SNMs are used as substrates by the *E. coli* SAM decarboxylase and can bind to the DNA adenine methyltransferase enzyme under *in vitro* conditions. Our findings underscore the role of structural features outside the active-site, such as the flexible loop, in determining MAT nucleotide selectivity and indicate that the SAM nucleotide analogues are likely to participate in downstream cellular metabolism.

## Results

### MAT structures reveal substrate-dependent ordering of a flexible active-site loop

MAT catalyzes the reaction of ATP and methionine to produce SAM (**Figure 1A)**.^18^ To understand the structural diversity of MAT, we exhaustively examined MAT crystal structures (Enzyme Commission number EC 2.5.1.6) from the RCSB PDB and obtained 120 structures from 21 different organisms (4 archaea, 4 eukaryotes, and 13 bacteria) (**Table S1**). The comparative structural alignment with the *Ec*MAT (PDB: 7LOO) as a reference showed that bacterial and eukaryotic MATs display low Root Mean Square Deviation (RMSD) values in the range of 0.478 Å - 1.171 Å whereas the archaeal ones show high RMSD, 5.477 Å - 8.977 Å (**Figure S1A-E)**. However, the archaeal MATs have low RMSD (0.894 Å - 2.003 Å) among themselves **(Figure S1B, D, E, G)**. Our finding supports existing literature on the structural and sequence divergence of archaeal MAT from bacteria and eukaryotes (**Figure S1C-G**).^23^

**Figure 1:**
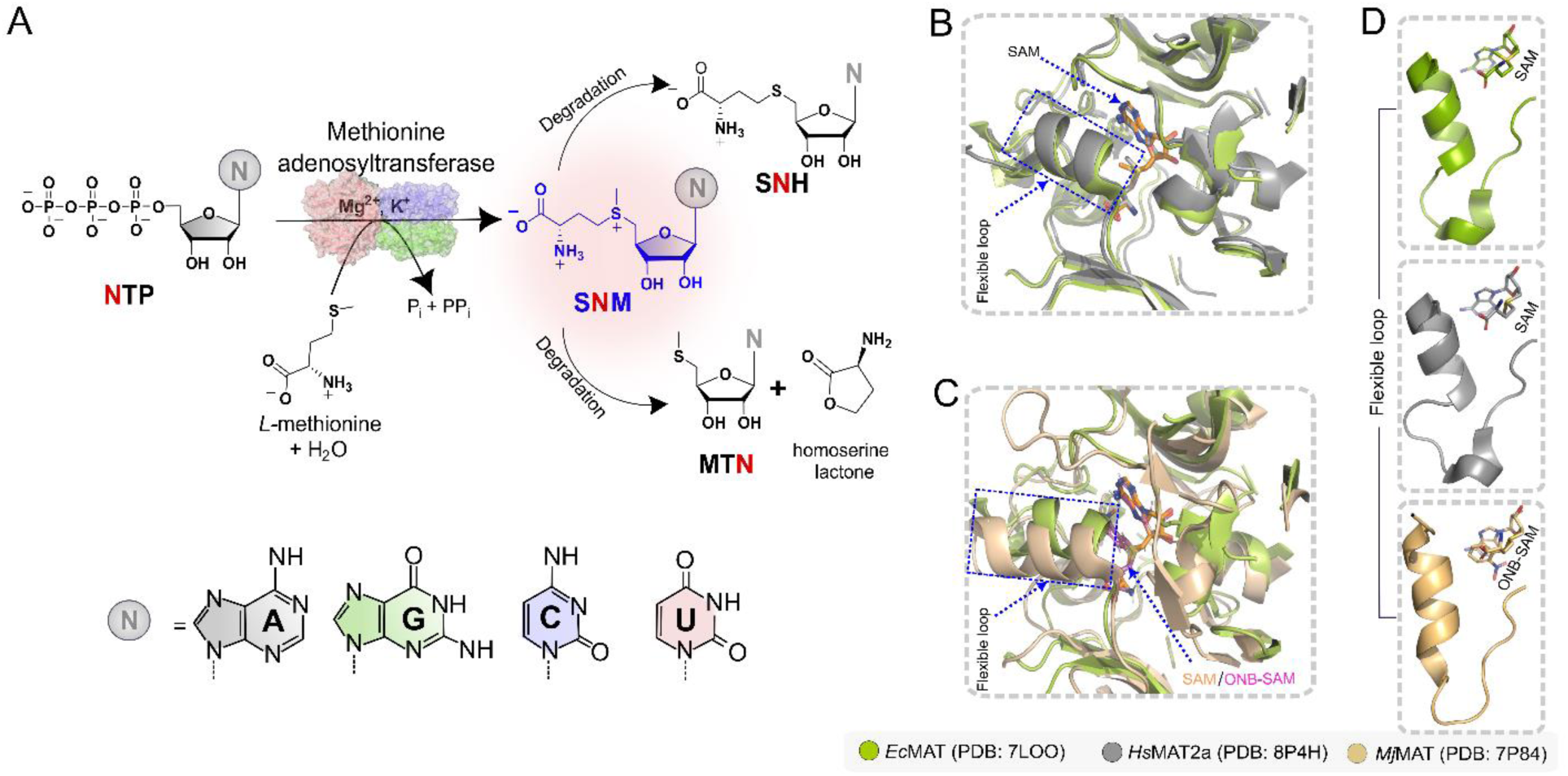
Enzymatic synthesis of nucleotide analogues of *S*-adenosyl-*L*-methionine using nucleotide-promiscuous methionine adenosyltransferase. **A**. The enzyme methionine adenosyltransferase (MAT) catalyzes the reaction between ATP and *L*-methionine to form *S*-adenosyl-*L*-methionine (SAM). NTP-promiscuous homologs of MAT can accept other NTPs as substrates to form the respective nucleobase analogues of SAM. The cartoon structure shown in the figure is that of *Escherichia coli* MAT (*Ec*MAT, PDB: 7LOO). **B**. Structural overlay of *Ec*MAT and *Homo sapiens* MAT2a (*Hs*MAT2a, PDB: 8P4H) shows that they are structurally very similar with RMSD of 0.661 Å. **C**. *Methanocaldococcus jannaschii* MAT (*Mj*MAT, PDB: 7P84) and *Ec*MAT are compared and they are structurally divergent with an RMSD of 6.66 Å. Crystal structures are visualised in PyMOL, residues within a radius of 1.5 nm around SAM or ONB-SAM are shown. **D**. The flexible loop regions of each MAT are shown with respect to the ligand bound.

Next, we examined all the MAT crystal structures for the presence of ligands bound in the active site. We found that in ligand-free structures, or in structures containing only one bound substrate i.e., ATP or methionine or their analogues, the loop is partially disordered or unresolved (**Figures S2A-E**). In contrast, if the MAT structure is co-crystallized with both substrates (methionine and an ATP analogue) or with the product SAM, this loop is in its folded state (**Figure S2F, G**). Interestingly, molecular dynamics simulations conducted previously have shown high root mean square fluctuation values for this loop in the apo form and when bound to only a nucleotide substrate, whereas it forms a structured helix on binding nucleotide and methionine or SAM.^15^ Also, the speculated role of the initial segment of the loop (residues 98-108) is to control substrate entry into the active site while a detailed study has identified the latter part (residues 108-122) of the *Ec*MAT flexible loop to modulate the efficiency of SAM formation from ATP and methionine.^20,24^ Based on existing literature and our comparative structural analysis, we hypothesized that this flexible loop might also be involved in the nucleotide selectivity of MAT. ^15,20,24^ As high-resolution crystal structures are available for *Ec*MAT (PDB ID: 7LOO), *Hs*MAT2a (PDB ID: 8P4H), and *Mj*MAT (PDB ID: 7P84), we used these three homologs for our next steps.^13,15,19^

### Nucleotide specificity of *Ec*MAT is unaltered by mutations in the methionine and adenine binding regions of the active site

*Ec*MAT is reported to show high selectivity for ATP as a substrate for reaction with methionine whereas *Mj*MAT and *Hs*MAT2a can utilize other nucleotides ^15,17,18^. First, using already reported assay conditions, we replicated the activity of *Ec*MAT and *Mj*MAT *in vitro* with ATP, GTP, CTP, and UTP.^17,18,25^ High performance liquid chromatography (HPLC) analysis confirmed *Mj*MAT uses ATP, GTP, CTP and UTP as substrates while *Ec*MAT primarily uses only ATP.^17,18^ As compared to *Ec*MAT, the activity of *Mj*MAT is 1.4 (± 0.2)-fold higher with ATP, 18 ( ± 0.5)-fold higher with CTP, 21.3 (± 0.4)-fold higher with GTP and 151.7 (±10.8)-fold higher with UTP, calculated for each reaction using the peak areas for SNM, and breakdown products *S*-nucleosyl-homocysteine (SNH) and 5’-methylthionucleoside (MTN) formed under previously reported reaction conditions **(Figure 2, Figure S3)**.^25^

**Figure 2:**
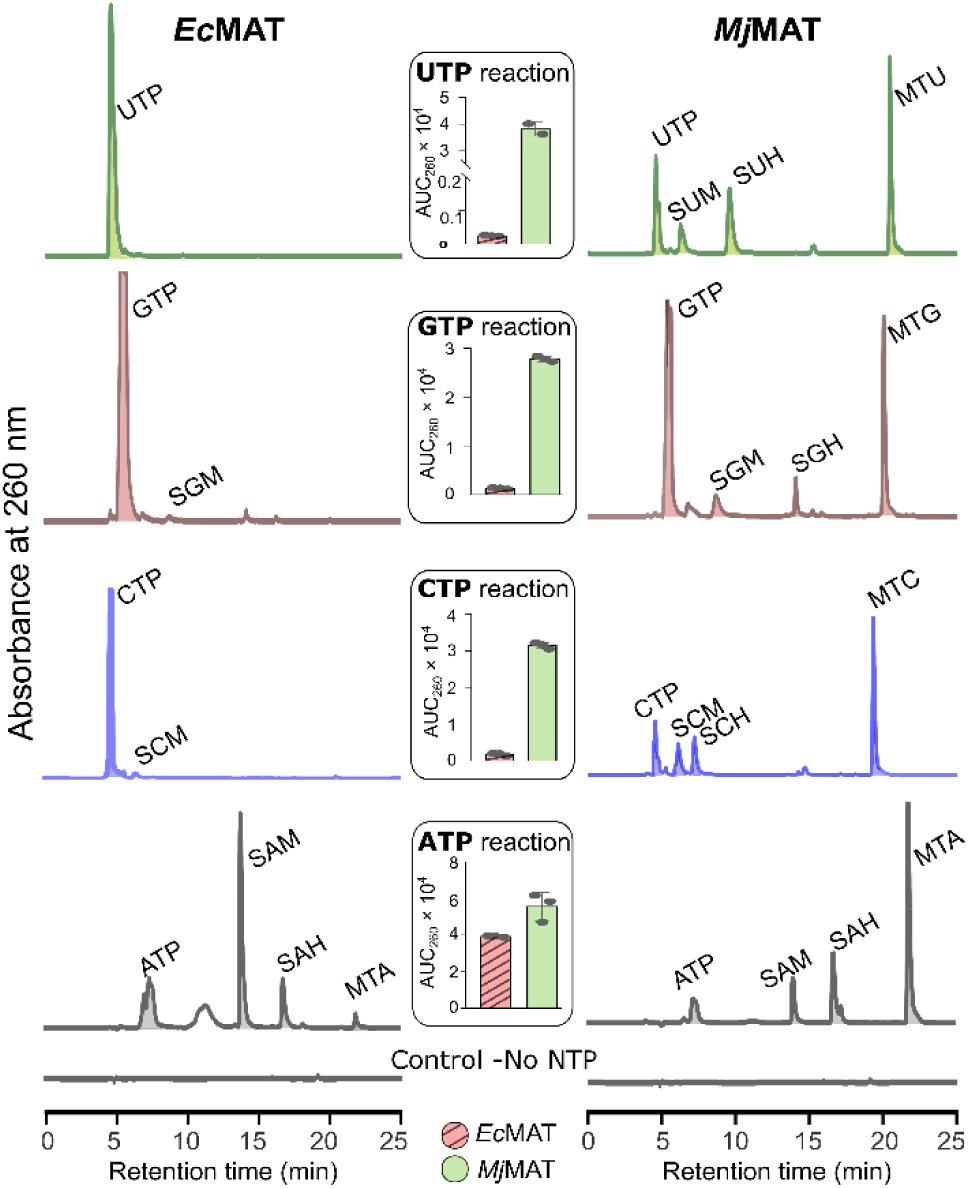
*Methanocaldococcus jannaschii* MAT (*Mj*MAT) exhibits NTP promiscuity, while *Escherichia coli* MAT (*Ec*MAT) shows ATP specificity. HPLC chromatograms of *Mj*MAT and *Ec*MAT reactions with ATP, GTP, CTP, and UTP. *Mj*MAT reactions show the formation of SNMs and their degradation products, for all four NTPs while *Ec*MAT shows product formation only with ATP. The bar graphs show the area under the curve (AUC) for all the product peaks (SNM, SNH and MTN) having absorption at 260 nm wavelength. Reactions are quenched after 12 hours and analyzed using HPLC.

To probe the differences in nucleotide utilization among the three MAT homologs, we compared their active site residues (**Figure 3A)**. The active sites of *Ec*MAT and *Hs*MAT2a are 95% identical in sequence and similar in structure (RMSD 0.478 Å) (**Figure 1B, S4A**).^15^ On the other hand, *Mj*MAT and *Ec*MAT diverge in terms of residues and overall structure (RMSD of 7.462 Å) (**Figures 1C, S4A).** In contrast, the active sites for *Ec*MAT, *Mj*MAT and *Hs*MAT2a are structurally similar and the orientation of product bound in the crystal structures are comparable (**Figure S4B-E)**.^15^ Additionally, each of them displays a flexible loop around the active site (residue numbers *Ec*MAT: 98-118, *Mj*MAT: 142-162, *Hs*MAT2a: 112-132) which are structurally similar (RMSD of 0.814 Å and 2.415 Å for *Hs*MAT2a and *Mj*MAT loops with respect to *Ec*MAT loop respectively) (**Figure 1D**).

**Figure 3:**
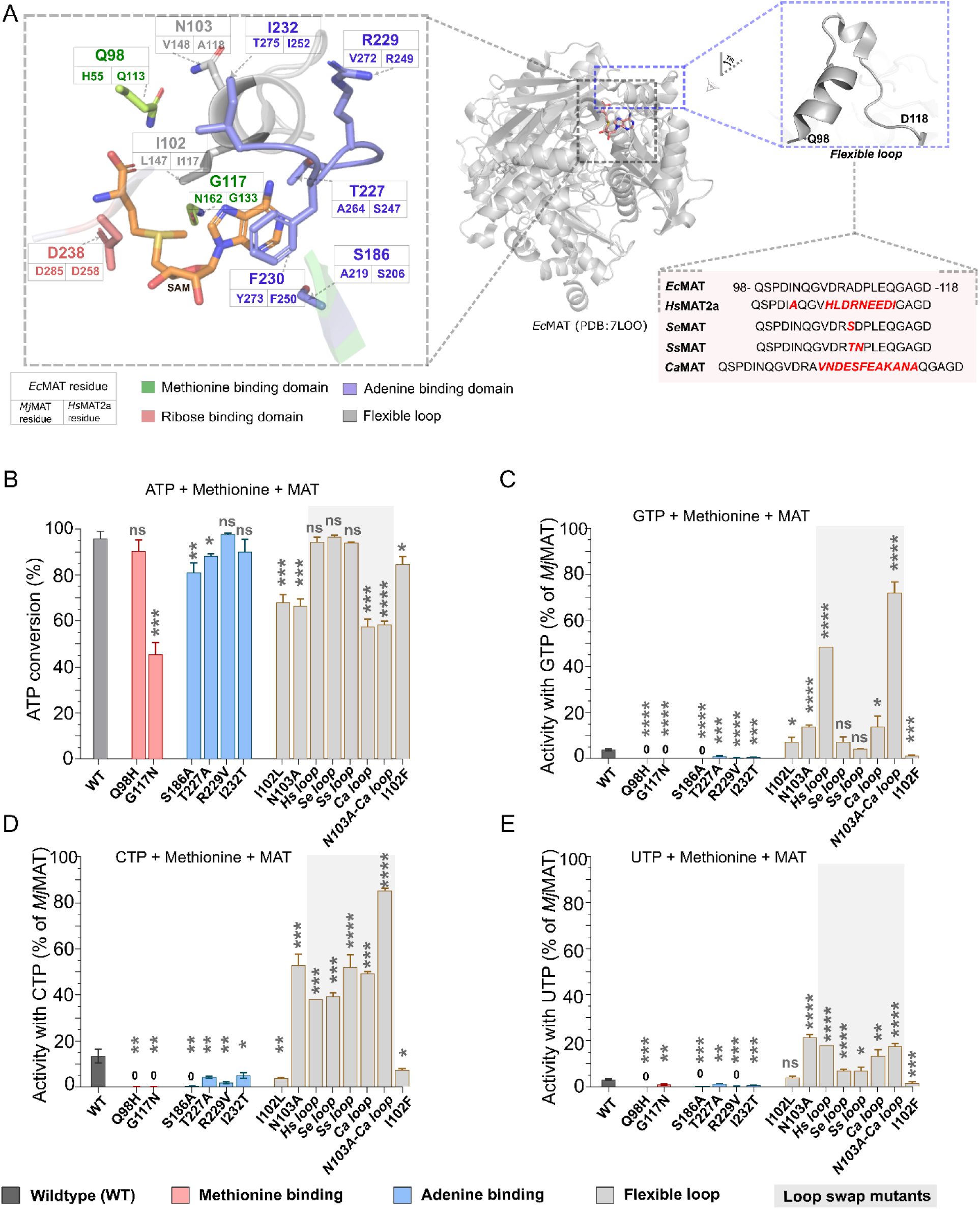
Rational design mutagenesis of *Ec*MAT for probing its nucleotide selectivity. **A**. The active site of SAM bound *Ec*MAT is shown. The residues in *Mj*MAT and *Hs*MAT2a are shown along with the corresponding residues of *Ec*MAT. The panel on the right side shows the flexible loop of *Ec*MAT. The amino acid sequence of the loop varies for MAT from different organisms. **B-E**. The activity of the mutants with all four NTPs is shown in the graphs. For ATP, the activity is shown as percentage conversion of ATP while for the other NTPs, the activity is shown with respect to *Mj*MAT as a reference. Datapoints are represented as the mean of three replicates +/− standard deviation, statistical test is two tailed unpaired Student’s t-test, p-values are shown as: ns (p>0.05), * (p<0.05), ** (p<0.01), *** (p<0.001), **** (p<0.0001). All comparisons are made with respect to wild-type *Ec*MAT.

To understand how the active site residues play a role in the nucleotide choice of *Ec*MAT, residues within a 5 Å radius of the bound SAM in the *Ec*MAT structure were considered for mutagenesis (**Figure 3A**). Point mutations Q98H, G117N in the methionine binding region, S186A, T227A, R229V, I232T in the adenine binding domain, and I102L in the flexible loop were proposed based on the residue comparison with *Mj*MAT, while N103A was inspired from *Hs*MAT2a flexible loop sequence. Additionally, based on the proximity of side chain of I102 with the adenine ring of ATP, we hypothesized that a phenylalanine in its place (I102F) might cause π-π stacking interaction with the aromatic purine and pyrimidine rings, thus allowing ATP and other NTPs to bind better. The ribose binding domains were identical, and the residues were conserved in each of the three MATs and so were not considered for mutagenesis.

All the point mutant proteins were cloned, purified, reactions with ATP, GTP, CTP and UTP were conducted and analyzed by HPLC.^18,25^ The activities of all the mutants with all four NTPs are shown (**Figure 3B-E, S5, table S2)**. All of the mutants retained >50% activity with ATP as compared to the wild type (**Figure 3B**). Even very significant changes like R229V, which is expected to disrupt a side-chain hydrogen-bonding interaction with the adenine base, did not substantially reduce ATP turnover. However, activity with noncognate NTPs decreased, indicating that this mutation preserves or strengthens ATP selectivity rather than broadening substrate scope. Strikingly, none of the other mutations in the residues that directly contact the adenine ring or methionine showed significant activity with GTP, CTP, and UTP and in fact mostly showed an overall drop in catalytic efficiency with respect to the wild-type *Ec*MAT **(Figure 3C-E)**. Overall, these results suggest that mutations in the adenine- and methionine-contacting residues tested here in *Ec*MAT are insufficient to broaden nucleotide specificity.

### Nucleotide specificity of *Ec*MAT is altered by mutations in the flexible loop region

Interestingly, I102L and N103A mutants, which are in the flexible loop region, showed a significant increase in activity with other NTPs **(Figure 3B-E)**. Although the activity with ATP for both the mutants were lowered to about 0.7-fold as compared to the wild-type, the N103A variant activity increased 3.6-fold, 3.9-fold, and 6.9-fold with GTP, CTP, and UTP, respectively while the I102L variant activity increased 1.9-fold and 1.2-fold with GTP and UTP, respectively and dropped 3.7-fold with CTP.

To investigate how the loop governs nucleotide specificity or promiscuity of MAT, we analyzed the composition of the 21 amino acid residues 98-118 in the *E. coli* MAT loop by comparing it with 5000 non-redundant MAT loop sequences and calculated the frequency of each amino acid at each position. The resulting position specific scoring matrix (PSSM) (**Figure S6A**) indicated that the five residues at the starting (residue numbers 98-102) and the five at the ending (residue number 114-118) are fully conserved, with variations in the middle. To understand the variations in loop length and sequence, we picked 20 distinct candidates from the 5000 non-redundant MAT sequences which had an overall loop sequence identity >30% with *E. coli* and >80% identity for the start and end sequences and loop lengths ranging from 21-37 owing to replacement/ insertions of amino acids (**Figure S6B**). Of these, we created the following four loop swap mutants - *Salmonella enterica* loop swap *(Se-*loop-*Ec*MAT), *Serratia symbiotica* loop swap (*Ss-*loop-*Ec*MAT), *Chryseobacteria arthrosphaerae* loop swap (*Ca-*loop-*Ec*MAT), and *Homo sapiens* loop swap (*Hs-*loop-*Ec*MAT) **(Figure 3A)**. All these loop swap mutants of *Ec*MAT showed increased activity with all four NTPs (**Figure 3C-E** grey bars, **S5**). In particular, the *Hs-*loop-*Ec*MAT mutant significantly gained activity with GTP (13-fold), CTP (3-fold) and UTP (6-fold), as well as retaining nearly 100% activity with ATP. Additionally, the *Ca*-loop-*Ec*MAT mutant showed high promiscuity, prompting us to create the N103A-*Ca*-loop-*Ec*MAT mutant, which shows a 19-fold, 6-fold and 4-fold enhancement of SGM, SCM, and SUM formation, respectively, as compared to the wild-type *Ec*MAT. Together, point-mutants and loop-swap data identify the flexible loop as a major determinant of MAT nucleotide selectivity.

### Production of nucleobase analogues of SAM

To test whether the engineered products could function as biochemical substrates or cofactors, we enzymatically synthesized SCM, SGM, and SUM by using the respective NTPs and the appropriate promiscuous *Ec*MAT mutants, and optimized pH conditions (**Figure S7**). We purified the SNMs using HPLC, confirmed their identities via Liquid Chromatography – Mass Spectrometry (LC-MS), and determined their UV-visible absorbance spectra (**Figure S8)**. The concentration of the SNMs was determined using the molar extinction coefficient of the respective NTPs (**Figure S9**). Being charged molecules with inherent instability, SNMs undergo degradation to form SNH and MTN analogues as shown (**Figure S3).** We purified and confirmed the identity of MTA, MTG, MTC and MTU by Nuclear Magnetic Resonance Spectroscopy (NMR) (**Figure S10)**.

### SNMs can partially rescue the growth of an *E. coli* SAM auxotroph

Having purified and characterized the SNMs, we wanted to understand whether they would be physiologically relevant for cellular metabolism. To address this question, the SAM auxotroph *E. coli* MOB1490 (a generous gift from Robert Blumenthal), created by introducing a SAM transporter from *Rickettsia prowazekii* followed by a knockout of *metK* (the gene coding for MAT), was used.^26,27^

*E. coli* MOB1490 can grow with either exogenous supplementation of SAM or with heterologous complementation of *metK* which encodes for a MAT protein (**Figure 4A**, **S11).**^27–29^ To see whether the SNMs support cellular metabolism and thus the growth of the SAM auxotroph, we fed these to MOB1490 cells. We conducted growth assays in LB medium supplemented with 200 μM SGM, SCM, or SUM, keeping SAM (200 μM and 2 μM) as positive controls, and observed that (i) none of the SNMs alone were able to support growth (**Figure S12A**) and (ii) final growth with 2 μM SAM is 3.6-fold lower than with 200 μM (**Figure 4B)**. Next, we repeated this assay with 200 μM SNMs + 2 μM SAM. As the purified SNMs were collected by HPLC using ammonium acetate as an eluent, an additional control, where only ammonium acetate was added to the cells along with 2 μM SAM was used. Interestingly, the 200 μM SNMs + 2 μM SAM combination showed a 1.3-1.8x fold increase in growth as compared to the 2 μM SAM (**Figure 4B**, top inset). Analysis using a sigmoidal growth model indicated that, as compared to only 2 μM SAM, the exponential growth rate increases with SCM + SAM, significantly slows down with SGM+SAM, and is not significantly altered with SUM + SAM **(Figure 4B**, bottom inset, **S12**). However, overall, the growth in SNM+SAM was consistently lower than when 200 μM SAM was provided. As the final growth of MOB1490 cells is a function of the transport as well as the cellular utilization of the SNMs, the differences in growth seen is a combination of both these factors. These results are consistent with the SNMs entering cells and supporting a subset of SAM-dependent functions, although transport and intracellular utilization cannot be distinguished from this assay alone.

**Figure 4:**
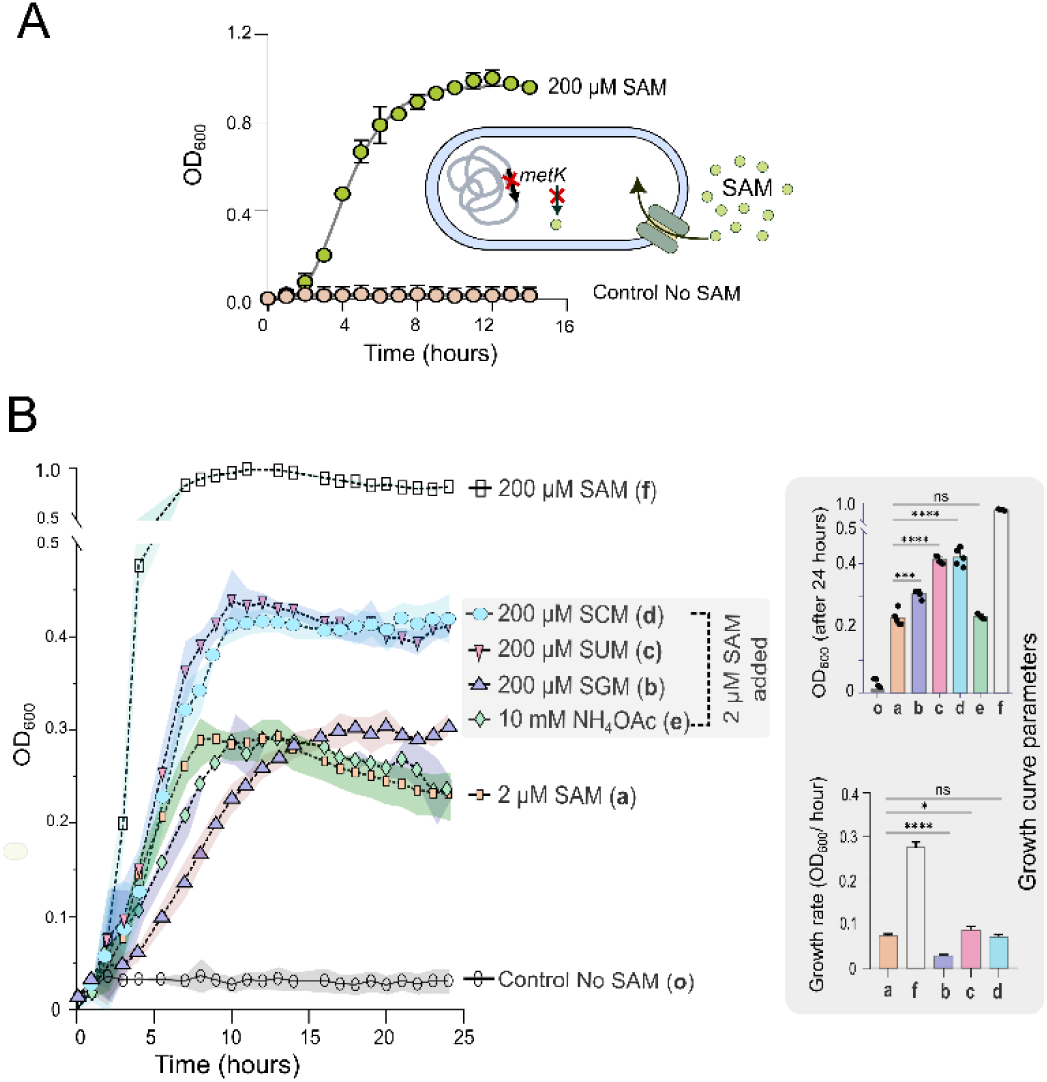
SAM nucleotide analogues partially rescue the growth of a SAM auxotroph. **A**. MOB1490 cells survive only if grown in media containing SAM. Because MOB1490 lacks metK, it requires exogenous SAM unless complemented; it also expresses a SAM transporter from Rickettsia prowazekii.^27^. **B.** MOB1490 is grown in LB media with additional supplementation of SAM and SNMs in different concentrations. Cells show increased growth with SAM+SNM (N=G/C/U) as compared to only SAM. NH_4_OAc supplementation control shows growth similar to SAM supplementation. The top inset bar graph indicates the end point OD_600_, while the bottom inset one shows the growth rates at the exponential phase. Datapoints are represented as the mean of 3-5 biological replicates +/− standard deviation, statistical test is two tailed unpaired students’ t-test, p-values are shown as: ns (p>0.05), * (p<0.05), ** (p<0.01), *** (p<0.001), **** (p<0.0001).

Because SNMs enhanced growth under SAM-limited conditions, we next tested whether representative *E. coli* SAM-utilizing enzymes could directly use these analogues – one, SAM decarboxylase that utilizes SAM as a substrate and two, DNA adenine methyltransferase that utilizes SAM as a methyl transfer cofactor.

### *E. coli* SAM decarboxylase accepts SGM, SCM and SUM as substrates

SAM decarboxylase, a crucial enzyme in *E. coli* for polyamine biosynthesis and an antimicrobial target, catalyzes the decarboxylation of SAM to give *S*-adenosyl-methioninamine (also called decarboxy-SAM or dcSAM) (**Figure 5A**).^30–33^

**Figure 5:**
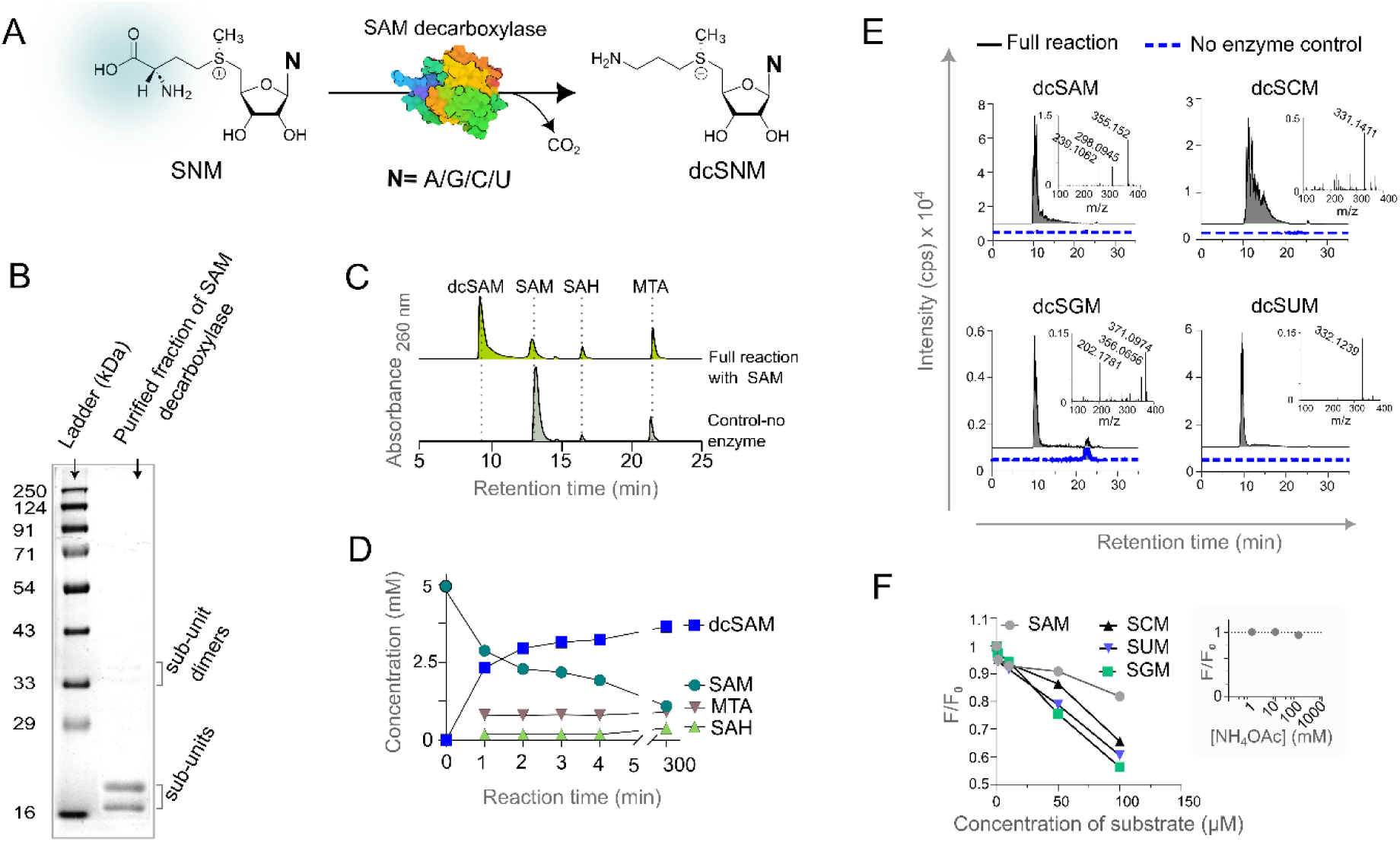
Nucleotide analogues of SAM as substrates for SAM decarboxylase. **A**. Reaction of SNM with SAM decarboxylase results in decarboxylated SNM (dcSNM). **B**. SAM decarboxylase is purified, using Ni-NTA column chromatography, as two distinct polypeptide sub-units bound together and analysed in an SDS-PAGE. **C**. A characteristic HPLC trace of the reaction mixture of SAM decarboxylase with SAM is shown. dcSAM elutes prior to SAM. The other two peaks are the degradation products of SAM, which are also present in the no enzyme control. **D**. The graph shows the concentration of the different species in the reaction of SAM decarboxylase with SAM. The molar extinction coefficients are assumed to be equal for SAM, dcSAM, SAH and MTA and concentrations are interpolated from standard curve of SAM (**Figure S13A**). As SAM can be enzymatically turned over to give dcSAM and non-enzymatically it degrades to SAH and MTA, the reaction mixture can only have these four components. **E**. The LC-MS chromatograms of the reaction mixtures of the SNMs with *E. coli* SAM decarboxylase are shown along with the corresponding mass fragmentation pattern of each dcSNM. The presence of the peak for dcSNM in the enzymatic reactions and their absence in the no enzyme control provides evidence of these SNMs being decarboxylated by SAM decarboxylase. **F**. Different concentrations of SNMs are titrated against SAM decarboxylase and the fluorescence is detected at 340 nm with excitation at 280 nm. The decrease in the signal ratio (F/F_0_) (fluorescence quenching) is consistent with enzyme–ligand interaction. As seen from the inset graph where signal does not drop with millimolar concentrations of ammonium acetate, the drop seen in case of SNMs is not the interference from the residual ammonium acetate remaining during collection of SNMs.

The His-tagged *E. coli* SAM decarboxylase enzyme was expressed and purified – two clear bands were observed on the gel, indicative of the post-translational internal cleavage characteristic of this enzyme (**Figure 5B**).^34^ To check its activity, a reaction was set up with commercially obtained SAM. The product decarboxy-SAM formed rapidly, confirmed by HPLC (**Figure 5C, 5D, S13A**). Next, to check whether SGM, SCM, and SUM can be decarboxylated, *E. coli* N103A MAT mutant was used to produce these molecules *in situ* and their reaction with SAM decarboxylase showed the formation of potential product peaks dcSGM, dcSCM, and dcSUM, although the conversion was poor **(Figure S13B-E).** To analyze the identity of these peaks, LC-MS analysis of the reaction was conducted with purified SAM nucleobase analogues. We observed m/z and a fragmentation pattern corresponding to dcSGM, dcSCM, and dcSUM along with dcSAM, establishing that the SNMs are indeed utilized as substrates by SAM decarboxylase (**Figure 5E**). Finally, we performed an intrinsic fluorescence quenching assay with different concentrations of enzymatically prepared purified SAM, SGM, SCM, and SUM and saw a decrease in normalized fluorescence intensity value (F/ F_o_) for the SNMs analogous to SAM, the native substrate **(Figure 5F)**.^25^ Together, the HPLC, LC–MS, fragmentation, and fluorescence-quenching data show that SGM, SCM, and SUM bind SAM decarboxylase and are converted to the corresponding decarboxylated products.

### *E. coli* DNA adenine methyltransferase can bind the SNMs but does not catalyze the methyl transfer

Next, we set out to probe SAM-utilizing enzymes involved in DNA methylation, a primary cellular function of SAM.^2^ We chose *E. coli* DNA adenine methyltransferase (DAM) which methylates the N^6^-adenine in 5’-G<u>A</u>TC-3’ of DNA (**Figure 6A**).^35,36^ The ASKA library plasmid pCA24N-*dam* was transformed, and the *E. coli* DAM protein was over-expressed and purified with some modifications of a previously reported method **(Figure 6B**).^36,37^

**Figure 6:**
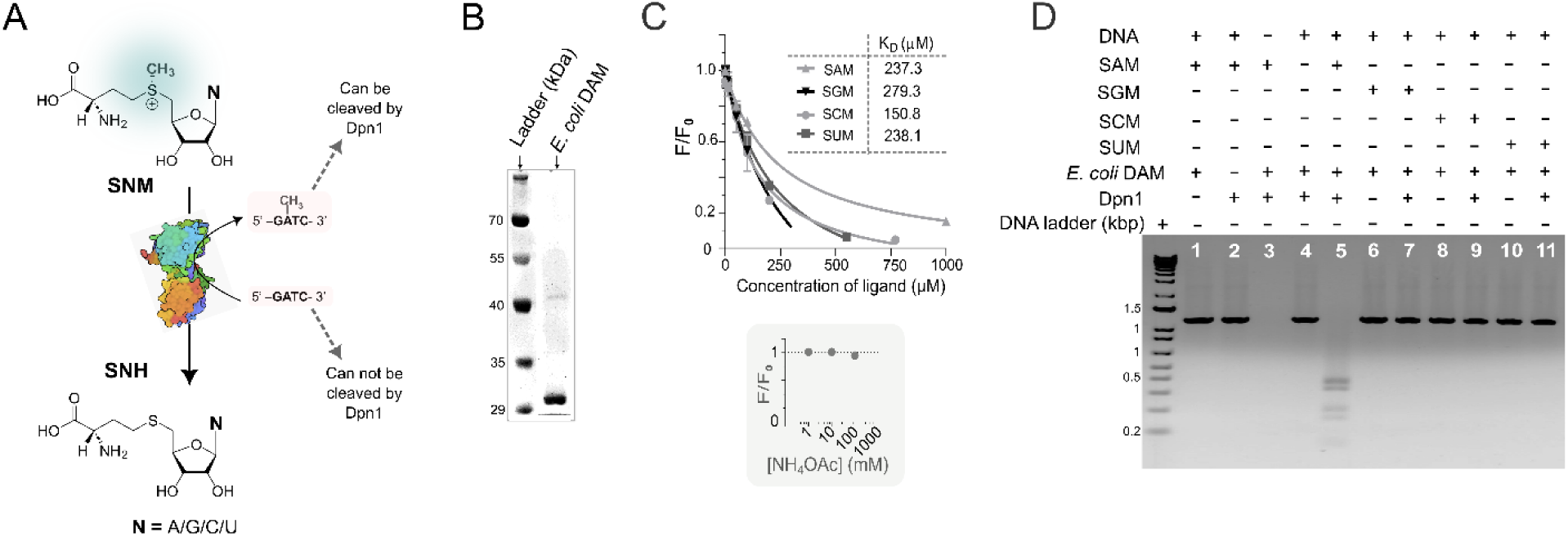
SAM nucleobase analogues bind *E. coli* DAM but do not support detectable DNA methylation. **A**. DNA adenine methyltransferase (DAM) catalyses the methyl group transfer from SAM (or SNMs) to the N^6^-adenine of DNA at 5’-GATC-3’ sites. SAH (or SNHs) is the other product in the reaction. Methylated DNA is cleaved by DpnI while the unmethylated DNA is not. **B**. *E. coli* DAM is purified by Ni-NTA affinity chromatography and analysed in an SDS-PAGE. It comes as a purified protein of size near to 32 kDa. **C**. Intrinsic fluorescence of *E. coli* DAM shows a decrease when SAM or the other SNMs are added, indicating the binding of these SNMs to the enzyme. As a control, addition of NH_4_OAc does not affect the fluorescence. **D**. The non-methylated DNA substrate is incubated with *E. coli* DAM and 5 μM SAM (or 50 μM of SGM, SCM or SUM) followed by DpnI digestion. Successful methylation renders the DNA susceptible to DpnI digestion, resulting in loss of the 1.1 kb substrate band and appearance of smaller cleavage products. Reaction mixtures lacking any one of the four components results in no reaction and no cleavage (lane 1-4). Lane 5 shows full cleavage of the DNA substrate; this is the reaction with SAM as the methyl donor. Similar reactions with SNMs (lanes 7,9 and 11) do not show any cleavage indicating unsuccessful methylation event. Data points are represented as the mean of two replicates +/− standard deviation.

To investigate whether the SNMs can bind to the *E. coli* DAM, we performed the intrinsic fluorescence quenching assay where binding of SAM causes a decrease in the intrinsic fluorescence intensity of the enzyme.^38^ A concentration-dependent decrease in fluorescence was observed when SAM or the other SNMs were added, indicating that they can bind to the enzyme **(Figure 6C).** Interestingly, SCM showed a lower apparent dissociation constant (K_D_) of 150.8 μM (as compared to SAM, K_D_ = 237.3 μM) under these assay conditions, indicating that SCM may bind DAM better than its native cellular substrate.

Next, we designed an activity assay for the *E. coli* DAM with some modifications from an earlier report.^39^ A 1.1 kb DNA fragment of non-methylated DNA substrate containing six 5’-GATC-3’ sites was incubated with SAM and *E. coli* DAM (sequence of the substrate DNA shown in **Figure S14**). Next, the DpnI restriction enzyme, which cleaves methylated and hemi-methylated DNA at the same 5’-GATC-3’ site, was added to initiate the reaction and analyzed by agarose gel.^40^ We observed that the reaction mixture containing DNA, SAM and *E. coli* DAM showed cleavage of the DNA on the addition of DpnI as expected **(Figure 6D, Lane 5)**. Next, we conducted the same reaction with SGM, SCM or SUM as the methylating agent. This time, we saw no significant DNA cleavage bands, indicating little or no detectable methylation of the substrate DNA **(Figure 6D, Lanes 6-11).** The results suggest that, despite being able to bind to *E. coli* DAM enzyme, the SAM analogues SGM, SCM, and SUM did not support detectable methyl transfer under these conditions. This may be attributed to an unfavourable orientation of the bound SNM inside the active site, hindering the catalysis.

Thus, for the two representative *E. coli* SAM-utilizing enzymes we tested, we conclude that the SCM, SGM, and SUM act as substrates for the SAM decarboxylase but cannot act as methylation cofactors for *E. coli* DAM even though they can bind to the enzyme. These enzyme-level results corroborate our previous observation that the SNMs alone cannot solely sustain the growth of the *E. coli* SAM auxotroph. (**Figure S12A**).

## Discussion

SAM is an essential cofactor used in methyl transfer, radical chemistry, polyamine biosynthesis, and several specialized metabolic reactions. Although the sulfonium center and methionine-derived portion of SAM are directly involved in many of these reactions, the adenine moiety is generally regarded as a recognition element. Whether SAM analogues with naturally occurring nucleobases, namely guanine, cytosine or uracil, can be formed within cells and/ or used as cofactors to sustain metabolism is not known, with only one example of MTG (a degradation product of SGM) being found in a human liver cancer cell line.^15^ Interestingly, MAT homologs have been reported to display diverse substrate scope *in vitro.*^15^ In this study, we investigated these two linked problems: what determines nucleotide selectivity in MAT, and whether SAM nucleobase analogues can function as cellular cofactors.

Our structural comparison of MAT homologs indicates that a flexible loop near the substrate-binding site undergoes ligand-dependent ordering. In ligand-free structures, or in structures containing only one bound substrate, this loop is often unresolved or disordered. In contrast, product-bound or ternary-complex structures show the loop in an ordered helix-loop-helix-loop conformation (**Figure S2F-G**).^15,20^ Previous structural and molecular dynamics studies have implicated this region in substrate entry and catalytic efficiency. Our data extend its role by showing that the same loop also contributes to nucleotide substrate recognition and thus nucleotide promiscuity.

By comparing the crystal structures and sequences of *Ec*MAT, *Mj*MAT and *Hs*MAT2a, we designed point mutants in the adenine binding, methionine binding and the flexible loop region. Although adenine binding and methionine binding domain mutants showed very little change in the nucleotide selectivity of *Ec*MAT, the flexible loop mutants N103A and I102L showed increased activity towards GTP, CTP and UTP. Interestingly, both these residues lie in the initial section of the helix of the flexible loop. A detailed analysis of the *Ec*MAT 21-amino acid long flexible loop sequence showed that the first and last five residues are highly conserved in MAT sequences. We identified 20 distinct loop sequences having >80% identity at the start and end sections and variations in the middle section and designed loop swap mutants *Se*-loop, *Ss*-loop, *Hs*-loop and *Ca*-loop *Ec*MAT where the residues far from the nucleobase were different. We observed that the loop swap mutants showed increased promiscuity with GTP, CTP and UTP. Notably, *Se*-loop (A109S), *Ss*-loop (A109S-D110N) and *Ca*-loop mutations are in the linker loop and the short helix of the flexible loop. This observation points towards the hypothesis that the two regions of the loop- the helix close to the nucleobase and the linker loop-helix away from the nucleobase, both are important for the nucleotide choice of *Ec*MAT. This hypothesis is strengthened by the observation that the N103A mutant of the *Ca*-loop *Ec*MAT, encompassing both the regions of the flexible loop, shows increased nucleotide promiscuity. Taken together, these results showed that, in addition to the conformational orientation of the different nucleotides in the active site, this flexible loop may act as a nucleotide ‘gatekeeper’, thus controlling the nucleotide promiscuity of MAT enzymes.^15^ An alanine-scanning or site-saturation mutagenesis of the gatekeeper loop might help to further refine the identities of the residues critically determining the catalysis or the conformational dynamics of nucleotides, by interacting with second or third shell of the active site loop and thus determining the choice of nucleotides in MAT enzymes.^15^

We next asked whether the products generated by nucleotide-promiscuous MAT variants could function beyond *in vitro* synthesis. Purified SGM, SCM, and SUM did not support growth of the *E. coli* SAM auxotroph MOB1490 when provided alone. However, when supplied together with a limiting concentration of SAM, the analogues increased final growth relative to limiting SAM alone. This result suggests that SAM nucleobase analogues can support a subset of SAM-dependent cellular functions, but cannot replace the full biochemical role of SAM. Because the growth phenotype reflects both transport and intracellular utilization, this experiment does not by itself distinguish whether differences among SGM, SCM, and SUM arise from uptake, stability, enzyme compatibility, or downstream metabolism.

To investigate the possible cellular utilization of SAM analogues within cells, we analyzed reactions with two representative SAM-utilizing *E. coli* enzymes - SAM decarboxylase and DAM. SAM decarboxylase accepted SGM, SCM, and SUM as substrates, producing the corresponding decarboxylated products. In contrast, DAM bound the analogues but did not support detectable methyl transfer to DNA under the conditions tested. These results indicate that the SAM nucleobase analogues can be treated as potential cofactors for some of the biochemical pathways inside the cell. The DAM result is especially important because it separates binding from catalysis. The fluorescence-quenching data indicate that the analogues can interact with the enzyme, but the DpnI-based methylation assay shows that binding alone is insufficient for methyl transfer. This likely reflects altered positioning of the cofactor within the active site. Thus, replacing adenine with guanine, cytosine, or uracil can preserve some recognition features while disrupting the precise geometry required for catalysis, putting forth the possibility of using these SNM analogues as inhibitors for SAM-binding/-utilizing enzymes. Further, whether these analogues can be biochemically synthesized in cells remains a question to be investigated.

Cofactor nucleotide analogs have also been designed for flavin adenine dinucleotide (FAD) and nicotinamide adenine dinucleotide (NAD), with exciting applications emerging from the cellular production of FAD and NAD nucleobase analogues.^41–43^ In that broader context, this work shows that SAM cofactor identity can also be diversified by engineering the nucleotide selectivity of its biosynthetic enzyme. More specifically, it identifies the MAT flexible loop as a tunable structural element for generating SAM nucleobase analogues and demonstrates that these analogues can participate selectively in SAM-dependent biology. This provides a foundation for SAM analogue production *in vitro* and within cells and developing semi-orthogonal cofactor systems for synthetic biology.

## Conclusion

In this work, we showed that a flexible gatekeeper loop contributes to the nucleotide selectivity of methionine adenosyltransferase (MAT). We engineered *Ec*MAT, which is selective for ATP as the nucleotide substrate and created variants that could also accept GTP, CTP and UTP as nucleotide substrates. By swapping the *Ec*MAT flexible loop with the corresponding loop sequences from different organisms, we obtained loop swap mutants which showed promiscuity for all four nucleotides. Further, we addressed a fundamental question whether SNMs, the nucleobase analogues of SAM, show cofactor-like function by first demonstrating that these SNMs can partially rescue the growth of an *E. coli* SAM auxotroph and that the SNMs can be substrates for *E. coli* SAM decarboxylase and can bind to *E. coli* DNA adenine methyltransferase but cannot catalyze the methyl transfer *in vitro*. This study not only provides a framework to investigate how the SAM nucleobase contributes to cofactor recognition and enzyme compatibility but also provides a foundation for engineering semi-orthogonal SAM cofactor systems.

## MATERIALS AND METHODS

### Materials

All primers, strains and plasmids used in this study are listed in Supplementary Information Tables S3-S5. The pET15b (WT *EcmetK*) construct was cloned in our lab, as previously reported.^25^ This vector produces N-6X-His-*Ec*MAT using the T7-lac expression system. *Escherichia coli* BL21 (DE3) strain containing pET19b (WT *MjmetK*) (that expresses N-10X-His-*Mj*MAT) was a gift from Prof. Squire J. Booker at Pennsylvania State University. *E. coli* MOB1490, a SAM auxotrophic strain, was kindly provided by Prof. Robert Blumenthal at the University of Toledo.^28^ It contains Δ*metK* (*metK*::Kan^R^), plasmid pMW1402 (Amp^R^) harbouring the SAM transporter from *Rickettsia prowazekii*^26,27^, and pMW1484 (Rif^R^), a pBAD-derived plasmid encoding *rpoB*. MOB1490 cells containing pUC57 (WT *EcmetK*) were also provided by Dr. Blumenthal, from which the pUC57 vector with *EcmetK* was extracted. *E. coli* K-12 AG1 strain with plasmids pCA24N-dam or pCA24N-speD, that expresses DNA adenine methyltransferase or SAM decarboxylase, respectively, were obtained from ASKA library.^37^

Media components were purchased from Hi-Media. Antibiotics were obtained from TCI chemicals and Sigma-Aldrich. 100 mM NTP solutions were procured from Jena Biosciences. L-methionine was obtained from TCI Chemicals. L-arabinose was obtained from SRL Chemicals. All other chemicals used were from Sigma-Aldrich (Merck). Protein ladders (10 to 250 kDa) were purchased from Genetix and ThermoFischer Scientific. DNA ladders (250 – 10,000 bp) were procured from NextGen. High-fidelity PrimeSTAR GXL Polymerase and DpnI were purchased from DSS TaKaRa Bio USA. Plasmid and DNA purification kits were obtained from Qiagen, Agilent, and Promega. Solvents for HPLC were procured from SD Fine Chemicals, Rankem, and Sigma-Aldrich. LC-MS grade methanol was purchased from J. T. Baker.

### Methods

#### Bioinformatic studies on the flexible loop

A protein BLAST (**B**asic **L**ocal **A**lignment **S**earch **T**ool available at https://blast.ncbi.nlm.nih.gov/Blast.cgilink) search was performed on the FASTA sequence of *Ec*MAT.^44^ The maximum number of search results was increased to 5000. A multiple sequence alignment of all these 5001 sequences was done using MUSCLE (**MU**ltiple **S**equence **C**omparison by **L**og-**E**xpectation available at https://www.ebi.ac.uk/jdispatcher/msa/muscle) and the loop sequences were sorted based on sequence identity with *Ec*MAT loop. Using a Bio Python module, the frequency of each amino acid in each of the positions of the loop (residues 98-118 in *Ec*MAT) were counted and this resulted in the Position Specific Scoring Matrix (PSSM). Also, 20 different sequences of the flexible loop were identified, and they were sorted based on their sequence identity with the corresponding loop in *Ec*MAT.

#### Restriction-free site directed mutagenesis on *Ec*MAT

Q98H, G117N, I232T, *Se-l*oop, *Ss-*loop and *Ca-*loop *Ec*MAT mutations were generated using megaprimer-based site-directed mutagenesis on pET15b-*EcmetK.*^45^ R229V, N103A, I102F, *Hs-*loop, I102L, T227A, and S186A mutations were introduced via Quick-Change site-directed mutagenesis.^46^ The successful introduction of the mutation in the plasmid is confirmed via Sanger sequencing. The pET15b-derived plasmids were sequenced using the T7 promoter and terminator.

#### Purification of *Ec*MAT, DAM methylase and SAM decarboxylase

The purification of all the proteins was done using slight modifications of previously reported protocols.^25,36^ The detailed methods are provided in supplementary information.

#### Activity assay of MAT

To check the activity of *Ec*MAT, enzymatic reactions are set up at 37 °C. Reaction mixture contains 50 mM HEPES buffer (pH 8.0), 50 mM MgCl_2_, 20 mM KCl, 15 mM *L*-methionine, 10 mM NTP and 20 μM enzyme. After 24 hours, reactions are quenched by adding 9% formic acid followed by centrifugation at 14000 rpm for 20-40 minutes to precipitate the enzyme. The supernatant is then injected into an analytic reverse phase column in HPLC. For bulk enzymatic synthesis of SNMs, HEPES buffer at pH 6.0 is used and reactions are quenched after 15-16 hours. The assay for *Mj*MAT uses the same reaction components, but the reaction is set at 55 °C.

#### High Performance Liquid Chromatography

The reverse phase HPLC of reaction mixtures was performed on an Agilent 1260 Infinity II UPLC system paired with a UV-Vis DAD detector, using an Agilent Zorbax Eclipse Plus C18 5μ 250mm x 4.60mm column. The solvent system comprises 10 mM pH 6.5 ammonium acetate (solvent A) and methanol (solvent B). The mobile phase gradient used was: 0-2 min 100% A, 2-15 min 100% A to 70% B, 15-20 min 70% B, 20-20.5 min 70% B to 100% B, 20.5-25 min 100% B, 25.0-25.5 min 100% B to 100% A, 25.5- 35 min 100% A. The chromatograms were recorded at 260 nm, as all four nucleobases strongly absorb at that wavelength. The HPLC traces were analyzed using Open Lab CDS and plotted using GraphPad Prism.

#### Liquid-Chromatography Mass Spectrometry (LC-MS)

Analysis of reaction mixtures and cell lysates using LC-MS was performed on ScieX X500R-QTOF mass spectrometer system attached to Exion-LC series UHPLC. The LC method is identical to the HPLC method used. The MS method is as follows: Electrospray ionization (ESI) was executed in the positive mode with +4.5 kV ion spray voltage, medium de-clustering potential (80V), and low collision energy (5V) at 400 °C. SNMs and their degradation products were identified using a quadrupole time of flight (QTOF) detector followed by targeted MRM-HR analysis using the following precursor and fragment ion masses: ATP (508.0, 508.0030), GTP (524, 523.9979), CTP (484, 483.9918), UTP (484.1, 484.0758), methionine (150.1, 150.0583), SAM (399.1, 399.1445), SAH (385.1, 385.1215), MTA (298.1, 298.0968), SGM (415.1, 415.1416), SGH (400.1, 400.1086), MTG (314.1, 314.0918), SCM (375.1, 375.1333), SCH (361.1103), MTC (274.1, 274.0856), SUM (376.1, 376.1173), SUH (362.1, 362.0943), MTU (275.1, 275.0696), decarboxy SAM (355.2, 355.1547), decarboxy SGM (371.1, 371.1496), decarboxy SCM (331.1, 331.1435) and decarboxy SUM (332.1, 332.1275). Data were processed on the ScieX-OS Explorer software to obtain extracted ion chromatograms (EIC), peak integrations, and the mass spectra of desired analytes.

#### Quantification of MAT nucleotide promiscuity

The formation of SAM from the reaction of MATs with ATP was quantified in the following manner. As the characteristic UV absorbance profile of SNM comes from the nucleobase which is common in all the significant degradation products of SNM, we can simply take the ratio of AUCs of leftover NTP and cumulative AUC_260_ of SNM, SNH and MTN to find the percentage of NTP remaining in the reaction mixture. A simple subtraction of this ratio from 1 followed by multiplication with 100 will give the percentage conversion of NTP. This works well for ATP. In the case of other SNMs, the turnover is low, leading to detector saturation for the NTP peak. Consequently, their formation could not be quantified using the above method. Instead, the activity of *Ec*MAT mutants with the other NTPs relative to WT *Mj*MAT is determined. This value was obtained by calculating the ratio of the AUC_260_ summation of the SNM, SNH, and MTN peaks from the *Ec*MAT mutant-NTP reaction trace to that obtained from the *Mj*MAT-NTP reaction trace.

#### Enzymatic synthesis, purification and quantitation of SNMs

To get pure SNMs from the enzymatic reaction of *Ec*MAT mutants with NTPs, reactions were set up in identical condition as mentioned above but with HEPES buffer at pH 6.0 (SNM degradation slows down at slightly acidic pH, **Figure S7**). After 15-16 hours, reaction mixtures were centrifuged at 14000 rpm for 30 minutes and then the supernatant is injected into HPLC. The flowthrough (post UV- detection) was collected at specific elution times corresponding to each SNM and then these fractions were concentrated in centrivap (speed vacuum). As the absorbance profile of SNM is characteristic of the nucleobase and as the molar extinction coefficient of SAM and ATP are experimentally determined to be nearly equal (**Figure S9A, B**), we assumed the molar extinction coefficients of other SNMs to be equal to that of the corresponding NTPs. So, we made a concentration vs absorbance standard curve for NTPs and then interpolated the concentration of SNMs from the corresponding standard curves.

The MTNs were collected using the same technique except that the reaction mixtures of *Mj*MAT at 55 °C are used, where MTNs are the major degradation product of the SNMs generated in the reaction mixture (**Figure 2**). The dried samples were dissolved in D_2_O, and NMR signals were acquired at a 400 MHz frequency NMR spectrometer.

#### MOB1490 feeding studies

MOB1490 cells were streaked from the glycerol stock (stored at −80 °C) on an LB agar plate containing 50 μM SAM, 50 μg/mL ampicillin, 25 μg/mL kanamycin, 50 μg/mL rifampin, and 5 mM L-arabinose and kept for overnight at 37 °C. Single isolated healthy colonies were picked and inoculated into LB broth containing 50 μM SAM, 50 μg/mL ampicillin, 25 μg/mL kanamycin, 50 μg/mL rifampin, and 5 mM L-arabinose and kept at 37 °C with 180 rpm shaking. Primary liquid culture was grown till OD_600_ of 0.5-0.7 (8-9 hours) and 1 mL from each culture corresponding to single colony was pelleted down at 5500 rpm for 2 mins. Cell pellet was resuspended in 1 mL LB broth, pelleted down and again resuspended in 0.5-1 mL LB broth. This was done to get rid of the carryover SAM from the primary culture. OD_600_ of the washed cultures was recorded in a 96 well microplate using the BMG Clariostar plate reader. Cells were inoculated in 200 μL media in each well (LB broth), with mentioned supplements, at a starting OD_600_ of 0.01 and allowed to grow for 24 hours at 37 °C with orbital shaking at 400 rpm. Datapoints were plotted and analyzed using GraphPad Prism software. For quantitative comparison and curve fitting, Gompertz model was used as previously described.^47^

#### Intrinsic Fluorescence assay for enzyme - substrate binding

To check the binding of SNMs with the enzymes of interest, intrinsic tryptophan fluorescence of the protein was used. The fluorescence is quenched upon ligand binding in the cases studied here. 1-10 μM enzyme in 100 mM Tris-Cl pH 8.0 was taken in a 96 well plate and different concentrations of SNMs were added to the wells. After 5 minutes of incubation at room temperature, fluorescence was measured. For fluorescence, excitation wavelength was 280 nm with a bandwidth of 5 nm in either side and the emission signal was collected at 360 nm with 10 nm bandwidth on either side. The graph of F/F_0_ was fitted using the equation.^25^ F=F_0_+((ΔF.[S])/([S]+K_D_)) where F is the intensity of the sample, F_0_ is the intensity of only enzyme control, ΔF is the change in fluorescence upon substrate binding, K_D_ is the apparent dissociation constant and [S] is the substrate concentration. From the best non-linear fit, we got the values of K_D_.

#### Activity assays of SAM Decarboxylase with SAM and SNMs

To check the activity of SAM Decarboxylase with SNMs, enzymatic reactions were set up at 37 °C. Reaction mixture contained 50 mM HEPES buffer (pH 7.4), 50 mM MgCl_2_, 20 mM KCl, 150 μM SNM and 5 μM enzyme. Reactions were quenched by adding 9% formic acid followed by centrifugation at 14000 rpm for 20-40 minutes to precipitate the enzyme. The supernatant was then injected into an analytic C-18 reverse phase column in HPLC and LC-MS. For *in-situ* generation of SNMs, *Ec*MAT-N103A reaction mixtures were setup as mentioned and *E. coli* SAM decarboxylase was added in the same reaction mixture after 24 hours.

#### Activity assays of DAM methylase with SAM and SNMs

To check the activity of DAM methylase, DNA substrate was prepared by amplification of a 1.1 kb fragment of DNA from pCA24N-*Dh*CobT plasmid. The sequence of the primers and the sequence of the DNA are listed (**Table S3, Figure S14)**. The DNA substrate has six *5’-<u>GATC</u>-3’* sites. The reaction was performed with slight modifications as described ^36^, in 100 mM Tris-Cl pH 8.0, 0.4 mg/mL BSA, 2.5 mM DTT, 5 or 50 μM SNM, 50 ng/ μL DNA substrate and 1 μM *Eco*DAM. The methylation reaction mixture was incubated at 37 °C for 30 minutes. After that DpnI and 10X T buffer were added to the reaction mixture in a 1:20 and 1:10 ratio, and the reactions were incubated for 1.5 hours at 37 °C. Reactions were then loaded in a 1.5 % agarose gel which was run for 40 minutes at a voltage of 90 V. DNA bands were visualized using ethidium bromide. Routine experiments were done with 20 μL volume of reaction mixtures.

## Acknowledgement

Financial support was provided by the Ministry of Science and Technology, Govt. of India –SERB Core Research Grant CRG/2019/003270, Anusandhan National Research Foundation Core Research Grant CRG/2023/002565, and partial support from the Higher Education Financing Agency for their Corporate Social Responsibility grant for this research to A.B.H. and for and KVPY-SX fellowship (by the Ministry of Science and Technology, India) to D.B.K. The authors acknowledge the DST fund for Improvement of S&T Infrastructure (SR/FST/LSII-043/2016) to IISER Pune Biology Dept for the Biological Mass Spectrometry Facility and the IISER Pune NMR and MS facilities and their managers. We sincerely acknowledge Professor Robert Blumenthal from University of Toledo for his kind gift of MOB1490 strain, and Shreyas Bhojane for his help in purifying SNMs from HPLC. We thank Shreya Mahato and Shradha Singh for critically reading the manuscript, Yamini Mathur for overall guidance and initial technical training for V. G. and A. B., and Aniket Vartak for his help with HPLC experiments. We acknowledge the use of ChatGPT Plus for checking for typographical, language, and citation errors.

