## Supplementary File for "Engineering a flexible loop in S-adenosyl-*L*-methionine synthetase enables production of SAM nucleobase analogues with selective biochemical and cellular activity"

[†] These authors contributed equally

**Present address:**

**Table S1: List of MAT crystal structures from different organisms.**

| Domain | Serial Number | Organism | Ligands bound | PDB ID | RMSD (Å) with 7LOO as reference |
|---|---|---|---|---|---|
| Bacteria | *1* | *Escherichia coli* | K, MG, PO4, POP, SAM | 7LOO | |
| | *2* | *Burkholderia pseudomallei* | * | 3IML | 0.817 |
| | *3* | *Mycobacterium marinum* | * | 3RV2 | 0.871 |
| | *4* | *Mycobacterium avium 104* | * | 3S82 | 0.865 |
| | *5* | *Mycobacterium tuberculosis* | * | 3TDE | 0.931 |
| | *6* | *Campylobacter jejuni RM1221* | * | 4LE5 | 1.152 |
| | *7* | *Cryptosporidium hominis* | 3PO, MG, SAM | 4ODJ | 0.569 |
| | *8* | *Thermus thermophilus HB27* | * | 5H9U | 0.781 |
| | *9* | *Neisseria gonorrhoeae FA 1090* | 3PO, AMP, MG, PO4, POP, SAM | 5T8S | 0.677 |
| | *10* | *Cryptosporidium parvum Iowa II* | CL, K, MG | 6C07 | 0.781 |
| | *11* | *Ureaplasma urealyticum serovar 7 str. ATCC 27819* | * | 6RJS | 0.982 |
| | *12* | *Lactiplantibacillus plantarum* | ANP, K, MG, PHE, PPK, SAM | 7R3B | 1.04 |
| | *13* | *Corynebacterium glutamicum ATCC 13032* | 3PO, ADN, GOL, K, MG, POP, SAM | 8JZG | 0.54 |
| Eukarya | *14* | *Entamoeba histolytica* | * | 3SO4 | 0.865 |
| | *15* | *Homo sapiens* (MAT2b)<br>*Homo sapiens* (MAT2a)<br>*Homo sapiens* (MAT1) | EDO, MG, PO4, SAM<br>CL, GOL, MG, SAM, UM6<br>* | 4KTT<br>8P4H<br>6SW5 | 0.478<br>0.466<br>0.915 |
| | *16* | *Arabidopsis thaliana* (MAT1)<br>*Arabidopsis thaliana* (MAT2) | CL, GOL, MG, MTA, SO4<br>MG, MPO, MXE, PGE, PGR, PO4 | 6VCY<br>6VCZ | 0.915<br>0.82 |
| | *17* | *Rattus norvegicus* | AMB, K, MG, SO4 | 1QM4 | 1.171 |
| Archaea | *18* | *Thermococcus kodakarensis* | * | 4L4Q | 8.357 |
| | *19* | *Saccharolobus solfataricus P2* | DPO, MG, PO4, SAM | 4L7I | 8.977 |
| | *20* | *Pyrococcus furiosus DSM 3638* | ACP, MDN, MG, PO4, SAM | 6S83 | 5.477 |
| | *21* | *Methanocaldococcus jannaschii* | * | 7P83 | 7.462 |

'*' indicates no ligand bound

**Table S1 legend**: Crystal structures with the highest resolution for MATs from different organisms used for comparative structural alignment analysis. The comprehensive list of PDB IDs of all other MAT crystal structures from the RCSB PDB database are mentioned here - 1FUG, 1MXA, 1MXB, 1MXC, 1O90, 1O92, 1O93, 1O9T, 1P7L, 1RG9, 1XRA, 1XRB, 1XRC, 2OBV, 2P02, 4HPV, 4K0B, 4KTV, 4L2Z, 4NDN, 4WS9, 5A19, 5A1G, 5A1I, 5T8T, 5UGH, 6FAJ, 6FBN, 6FBO, 6FBP, 6FCB, 6FCD, 6FWB,

6G6R, 6LTV, 6LTW, 6P9V, 6RK5, 6RK7, 6RKA, 6RKC, 6S81, 6SW6, 6VCW, 6VCX, 6VD0, 6VD1, 6VD2, 6WKB, 7BHR, 7BHS, 7BHT, 7BHU, 7BHV, 7BHW, 7BHX, 7KCC, 7KCE, 7KCF, 7KDA, 7KDB, 7L1A, 7LL3, 7LNH, 7LNN, 7LO2, 7LOW, 7LOZ, 7P82, 7P84, 7P8M, 7R2W, 7RW5, 7RW7, 7RWG, 7RWH, 7RXV, 7RXW, 7RXX, 8AXZ, 8JZH, 8JZI, 8OOG, 8P1T, 8P1V, 8P1W, 8QDY, 8QDZ, 8QE0, 8QE1, 8QE2, 8QE3, 8SWA, 8XAM, 8XAR, 8XB0.

**Table S2**: Fold change in activity of the mutants with respect to wild type, for all four NTPs.

| **Mutant *Ec*MAT** | **Fold change in activity with respect to *Ec*MAT wild type** | | | |
|---|---|---|---|---|
| | **ATP** | **GTP** | **CTP** | **UTP** |
| Wild type | 1.0 | 1.0 | 1.0 | 1.0 |
| Q98H | 1.0 | 0 | 0.1 | 0.1 |
| G117N | 0.5 | 0 | 0.1 | 0.4 |
| S186A | 0.9 | 0.1 | 0.1 | 0.2 |
| T227A | 1 | 0.3 | 0.4 | 0.5 |
| R229V | 1.1 | 0.1 | 0.2 | 0.1 |
| I232T | 1 | 0.2 | 0.4 | 0.3 |
| I102L | 0.8 | 2 | 0.3 | 1.3 |
| I102F | 0.9 | 0.5 | 0.6 | 0.2 |
| N103A | 0.7 | 3.7 | 3.9 | 7 |
| *Hs*-loop | 1 | 12.9 | 2.8 | 5.8 |
| *Se*-loop | 1.1 | 2 | 2.9 | 2.2 |
| *Ss*-loop | 1 | 1.2 | 3.9 | 2.2 |
| *Ca*-loop | 0.7 | 3.7 | 3.6 | 4.3 |
| N103A-*Ca*-loop | 0.7 | 19.2 | 6.3 | 5.7 |

**Table S3:** Primers used in the study:

| **Name of primer** | **Sequence (5' → 3')** |
|---|---|
| Q98H_MP_F | ATCGGCAAACATTCTCCTGACATCAAC |
| G117N_MP_F | GAACAGGGCGCGAACGACCAGGGTCTG |
| I232T_MP_R | ATTGGGCCACCGGTAACGAAACGA |
| R229V_QC_F | CGGTGTGTTCGTTATCGGTGGCCCAATGGG |
| R229V_QC_R | ATAACGAACACACCGGTCGGGTTGATGAAGAATTTGG |
| I102L_QC_F | CCTGACCTGAACCAGGGCGTTGAC |
| I102L_QC_R | CTGGTTCAGGTCAGGAGACTGTTTGCCGATAGCG |
| T227A_QC_F | CCCGGCGGGTCGTTTCGTTATCGGTGGCCC |

| | |
|---|---|
| T227A_QC_R | CGACCCGCCGGGTTGATGAAGAATTTGGTGGCAG |
| S186A_QC_F | GCTTGCGACTCAGCACTCTGAAGAGATCGACC |
| S186A_QC_R | CTGAGTCGCAAGCACGACAGCATCGATACCAAC |
| T7_promoter_F | TAATACGACTCACTATAGGG |
| T7_terminator_R | GCTAGTTATTGCTCAGCGG |
| I102F_QC_F | CCTGACTTTAACCAGGGCGTTGACCGTGCC |
| I102F_QC_R | CTGGTTAAAGTCAGGAGACTGTTTGCCGATAGCG |
| N103A_QC_F | GACATCGCGCAGGGCGTTGACCGTGCCGATCC |
| N103A_QC_R | GCCCTGCGCGATGTCAGGAGACTGTTTGCCG |
| *Hs* loop QC_F | GACATCGCGCAGGGCGTTCATCTGGACCGTGCCGATCCG CTGGAACAG |
| *Hs loop*_QC_R | GCACGGTCCAGATGAACGCCCTGCGCGATGTCAGGAGAC TGTTTGCCG |
| *Se* loop_MP_R | CTGTTCCAGCGGATCGCTACGGTCAACGCCC |
| *Ss* loop_MP_R | GTTCCAGCGGGTTGGTACGGTCAACG |
| *Ca* loop MP_F | GGGCGTTGACCGTGCCGTGAACGATGAAAGCTTTGAAGCG AAAGCGAACGCGCAGGGCGCGGGTGACC |
| N103A-*Ca loop_MP_F* | CCTGACATCGCGCAGGGCGTTGACCGTGCCGTGAACGAT GAAAGCTTTGAAGCGAAAGCGAACGCGCAGGGCGCGGG |
| CobT_F | TACGGATCCGGCCCTGAGGGCCGTGGATGACTTCTTAAGA CAGG |
| CobT_R | AAGCTTGGCTGCAGGTCGACCCTTAGTTTTTACCGCTGACT CC |

**Table S4:** Strains used in the study:

| Strain name | Genotype | Reference |
|---|---|---|
| MOB1490 | BW25113, Δ*metK (metK::Kan*$^R$*)* | [1] |
| *E. coli* DH5**α** | F⁻ φ80*lac*ZΔM15 Δ(*lac*ZYA-*arg*F) U169 *rec*A1 *end*A1 *hsd*R17($r_K^-$, $m_K^+$) *pho*A *sup*E44 λ⁻ *thi*-1 *gyr*A96 *rel*A1 | |
| *E. coli* BL21 (DE3) | *F⁻ompT hsdS*$_B$ *(*$r_B^-$*,* $m_B^-$*) gal dcm (DE3)* | [2] |

**Table S5:** Plasmids used in the study:

| Name | Gene expressed | Reference |
|---|---|---|
| pET19b-*MjmetK* | *MjmetK* | Gift from Prof. Squire J. Booker |
| pET15b-*EcmetK* | *EcmetK* wildtype | [3] |
| pET15b-*EcmetK-Se* loop | *EcmetK Sa Ec* loop swap | This study |
| pET15b-*EcmetK*-Q98H | *EcmetK* Q98H | This study |
| pET15b-*EcmetK*-G117N | *EcmetK* G117N | This study |
| pET15b-*EcmetK*-R229V | *EcmetK* R229V | This study |
| pET15b-*EcmetK*-I102L | *EcmetK* I102L | This study |

| | | |
|---|---|---|
| pET15b-*EcmetK*-I232T | *EcmetK* I232T | This study |
| pET15b-*EcmetK*-S186A | *EcmetK* S186A | This study |
| pET15b-*EcmetK*-N103A-*Ca*-loop | *EcmetK* N103A-*Ca*-loop | This study |
| pET15b-*EcmetK*-N103A | *EcmetK* N103A | This study |
| pET15b-*EcmetK-Hs l*oop | *EcmetK-Hs l*oop | This study |
| pET15b-*EcmetK-Ca l*oop | *EcmetK-Ca l*oop | This study |
| pET15b-*EcmetK-Ss* loop | *EcmetK-Ss* loop | This study |
| pET15b-*EcmetK*- I102F | *EcmetK* I102F | This study |
| pUC57-*EcmetK* | *EcmetK* wild type | [1] |
| pCA24N- *dam* | *dam* | ASKA [4] |
| pCA24N- *speD* | *speD* | ASKA [4] |

### Supplementary material- figures

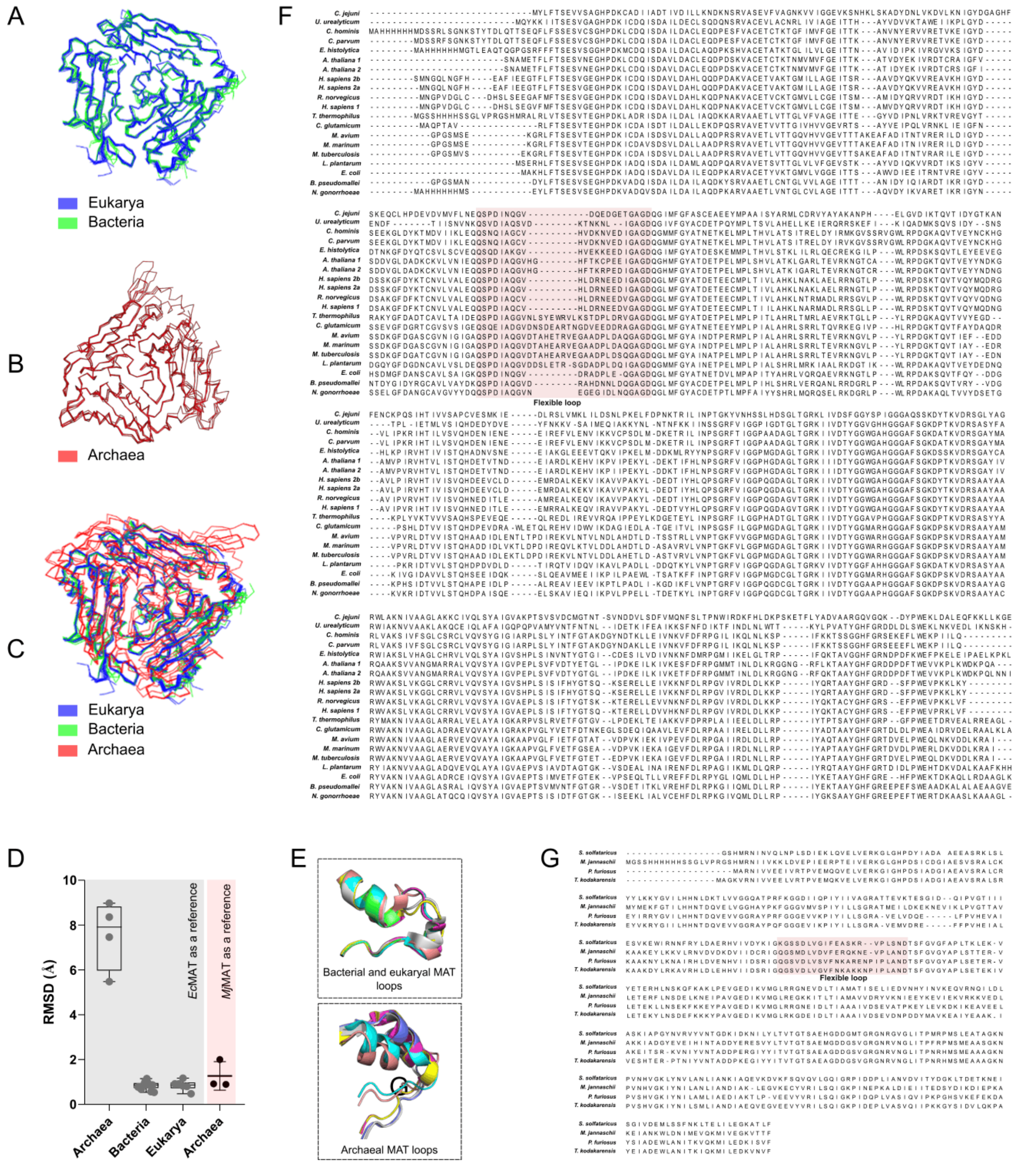


**Figure S1**. Structural and sequence alignment of MATs crystallized from different organisms. **A**. The bacterial and eukaryal MATs overlaid on *Ec*MAT (PDB:7LOO). They show very low RMSD values. **B**. Overlay of archaeal MATs on *Mj*MAT (PDB:7P84). Archaeal MATs align well within themselves, with less structural deviation. **C**. All the MAT crystal structures are overlaid on *Ec*MAT. The archaeal MATs show significant structural deviation with high RMSD values. **D**. The bar graph shows the RMSD values of MATs (with respect to *Ec*MAT or *Mj*MAT) from the three domains of life. The archaeal MATs are structurally divergent from the rest. **E**. The flexible loop sequences for the

SAM bound structure of MAT from different organisms are overlaid and shown. **F.** Alignment of 20 MAT sequences from 17 different bacterial and eukaryotic sequences. **G**. Alignment of 4 MAT sequences from archaeal organisms. The flexible loop sequences are highlighted in light red.

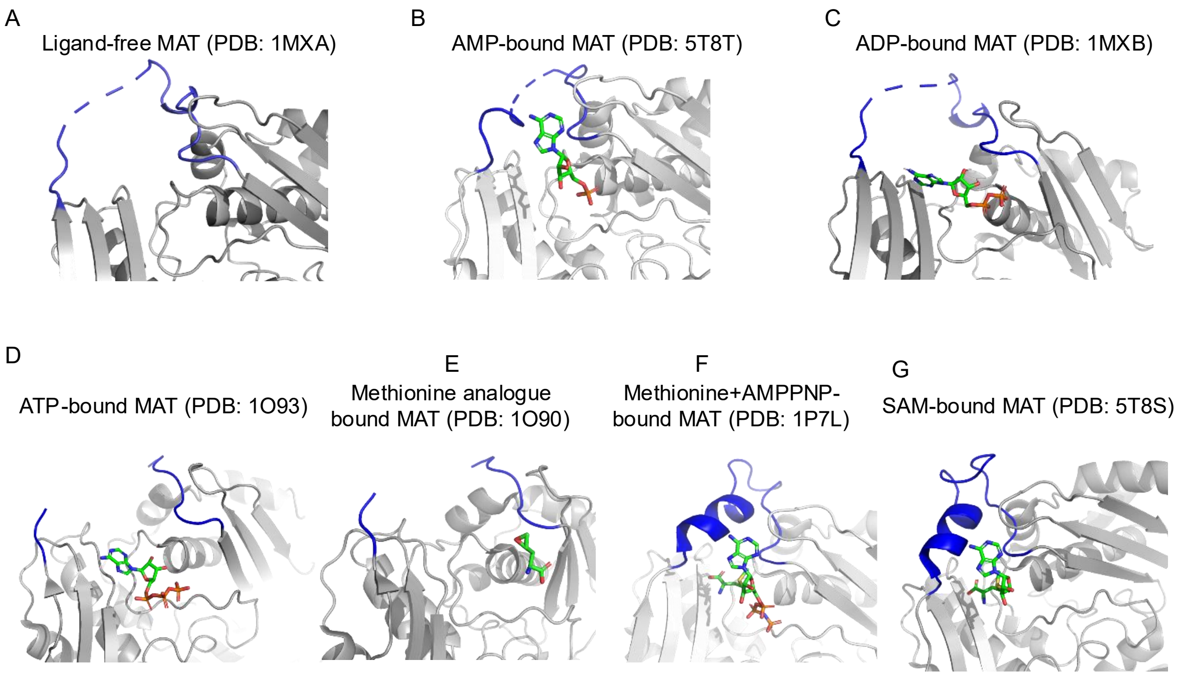


**Figure S2**. The flexible loop (blue color) is shown in different representative crystal structures of MAT in ligand-free (**A**) or different ligand bound states (**B-G)**. The loop electron density is unresolved in a crystal structure (PDB ID: 1O9T) although both ATP and methionine are bound, and the two ligands are not positioned correctly for an attack. This can be explained by the fact that the crystal was obtained by first crystallizing MAT with methionine and then soaking in ATP solution.

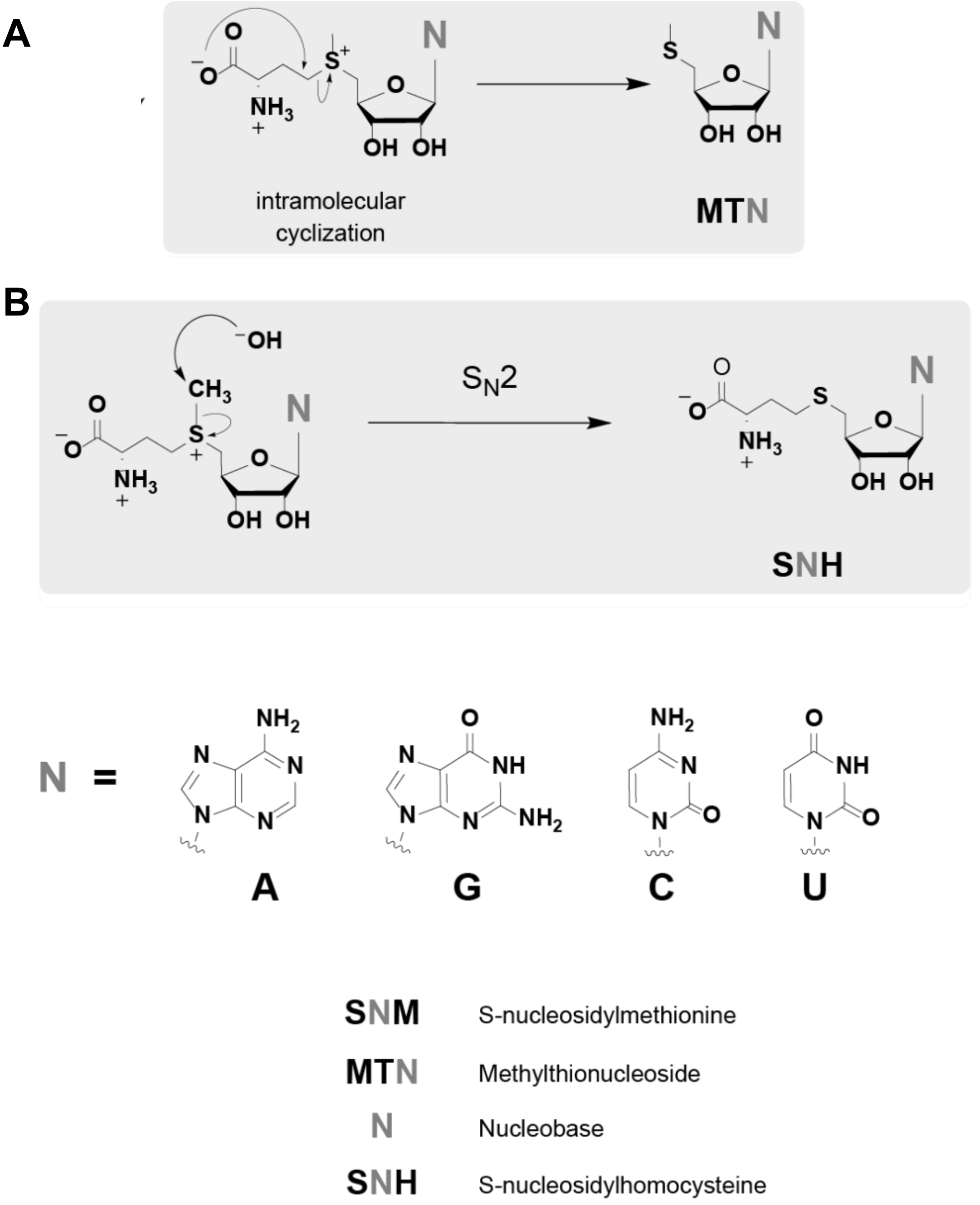


**Figure S3**: Mechanism of degradation of *S*-nucleosyl-*L*-methionine (SNM) to 5'-methylthionucleoside (MTN) **(A)** and *S*-nucleosyl homocysteine (SNH) **(B).**

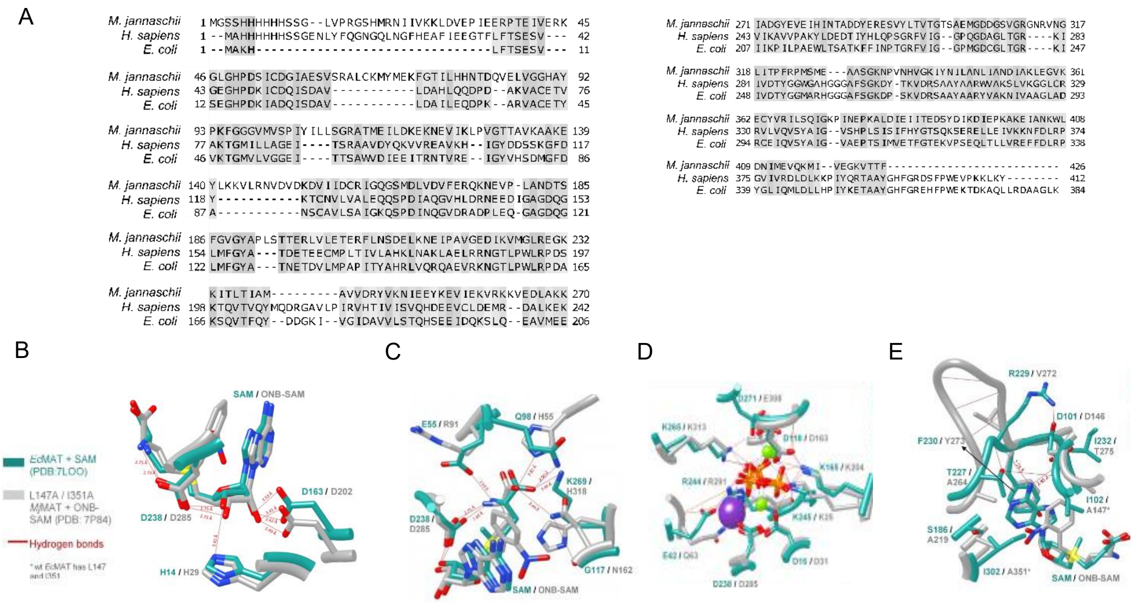


**Figure S4: A**. Alignment of the sequences of *Mj*MAT, *Ec*MAT and *Hs*MAT2a is shown. **B-E**. SAM and PPi bound *Ec*MAT (PDB:7LOO) and ortho-nitro benzyl SAM (ONB-SAM) bound *Mj*MAT (PDB:7P84) are represented using Chimera. Active site interactions within 5 Å shown with the ribose ring (**B**), methionine backbone **(C)**, triphosphate (**D**), and nucleobase (**E**)

#### Tab

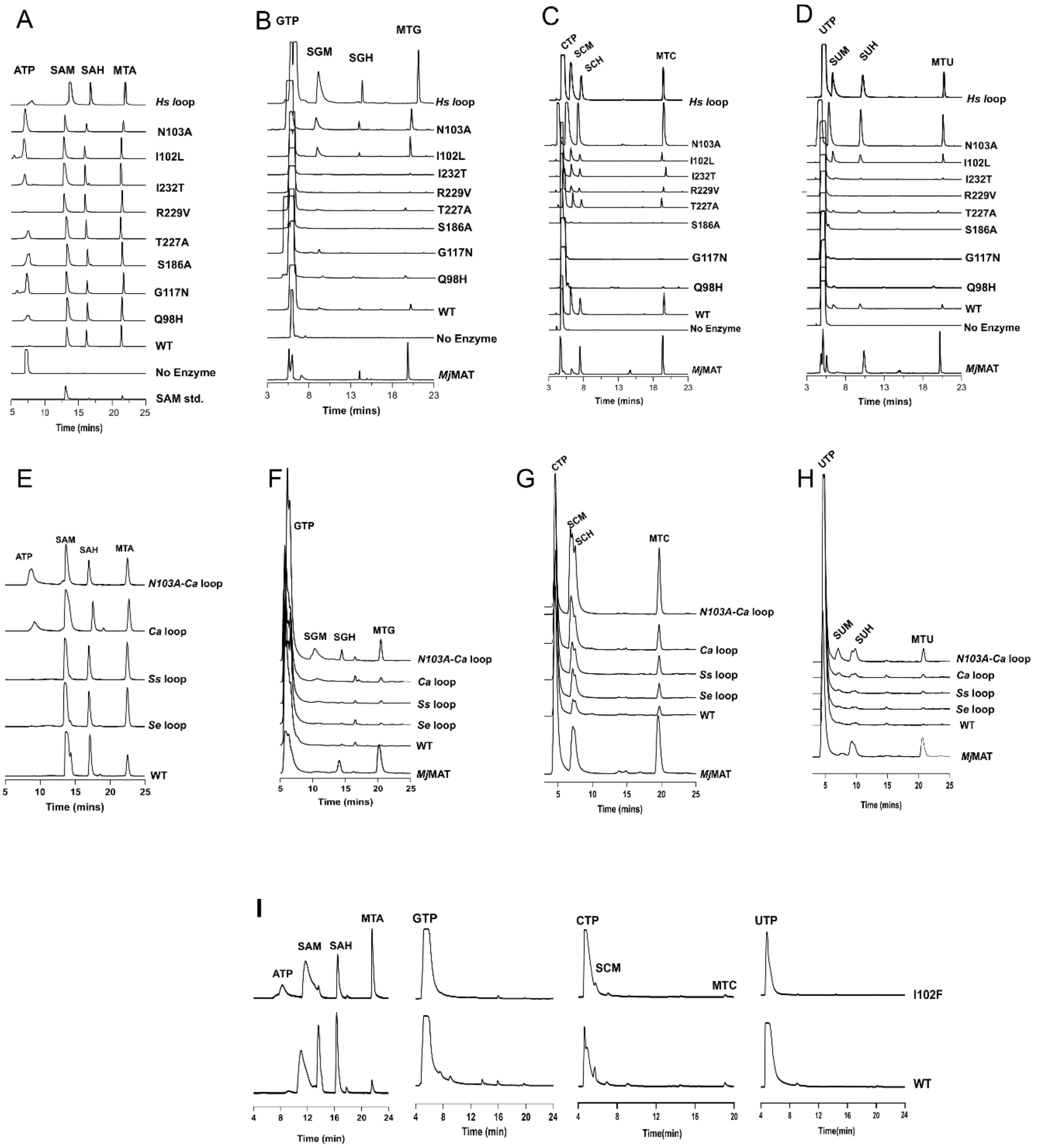


**Figure S5:** HPLC traces of the reaction mixture of all the mutants with ATP, GTP, CTP and UTP. Injection volumes are 10 µL (**A, E**), 55 µL (**B-D**), 20 µL (**I**) and 3 µL (**F-H**). The intensities across the different graphs are not to scale.

A

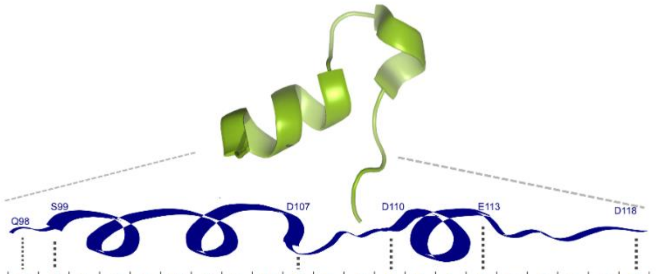


| | Q | S | P | D | I | N | Q | G | V | D | R | A | D | P | L | E | Q | G | A | G | D |
|---|---|---|---|---|---|---|---|---|---|---|---|---|---|---|---|---|---|---|---|---|---|
| A | 0 | 0 | 19 | 0 | 0 | 95 | 3.4 | 0.2 | 0 | 0 | 7.5 | 74 | 3.4 | 0.9 | 3.2 | 29 | 0.5 | 0 | 100 | 0 | 0 |
| C | 0 | 0 | 0 | 0 | 0 | 0 | 0 | 0 | 0 | 0 | 0.1 | 0 | 0 | 0 | 0 | 0 | 0 | 0 | 0 | 0 | 0 |
| D | 0 | 0 | 0.3 | 99 | 0 | 0.4 | 0.1 | 0 | 0 | 74 | 7.2 | 0.5 | 1.3 | 11 | 30 | 18 | 0.1 | 0 | 0 | 0 | 100 |
| E | 0 | 0 | 0.4 | 1.3 | 0 | 0 | 0.3 | 0 | 0 | 0 | 20 | 1.3 | 5.7 | 2.3 | 32 | 10 | 0 | 0 | 0 | 0 | 0 |
| F | 0 | 0 | 0 | 0 | 0 | 0 | 0 | 0 | 0 | 0 | 0 | 0.1 | 1.2 | 2.7 | 0.2 | 0 | 0.1 | 0 | 0 | 0 | 0 |
| G | 0 | 0 | 2.2 | 0 | 0 | 0.2 | 3.5 | 100 | 0 | 0.1 | 0.1 | 12 | 0.2 | 0.9 | 1.4 | 0.2 | 1 | 100 | 0 | 100 | 0 |
| H | 0 | 0 | 0.1 | 0 | 0 | 0 | 0 | 0 | 0 | 0 | 5.3 | 0 | 5.6 | 4.1 | 0.3 | 0 | 0 | 0 | 0 | 0 | 0 |
| I | 0 | 0 | 0 | 0 | 100 | 0 | 0.8 | 0 | 0.2 | 0 | 0 | 0.1 | 1.7 | 28 | 0.1 | 0.3 | 49 | 0 | 0 | 0 | 0 |
| K | 0 | 0 | 0.6 | 0 | 0 | 0.2 | 0 | 0 | 0 | 0.1 | 16 | 0.9 | 2.6 | 0.5 | 2.5 | 8.6 | 0.1 | 0 | 0 | 0 | 0 |
| L | 0 | 0 | 0.1 | 0 | 0 | 0 | 5.1 | 0 | 0 | 0.1 | 0.1 | 0 | 48 | 26 | 25 | 2.4 | 1.8 | 0 | 0 | 0 | 0 |
| M | 0 | 0 | 0 | 0 | 0 | 0 | 19 | 0 | 0 | 0 | 0.1 | 0 | 0.2 | 0.5 | 0.6 | 0.3 | 0.7 | 0 | 0 | 0 | 0 |
| N | 0 | 0 | 5.2 | 0.1 | 0 | 2.1 | 0 | 0 | 0 | 24 | 2.8 | 0.2 | 1 | 6.7 | 1 | 14 | 0.4 | 0 | 0 | 0 | 0 |
| P | 0 | 0 | 64 | 0 | 0 | 0 | 0 | 0 | 0 | 0 | 0.1 | 0 | 0.6 | 1.1 | 0.2 | 0 | 0 | 0 | 0 | 0 | 0 |
| Q | 100 | 0 | 2.2 | 0 | 0 | 0 | 65 | 0 | 0 | 0 | 12 | 0.3 | 1.4 | 0.2 | 0.3 | 3.2 | 38 | 0 | 0 | 0 | 0 |
| R | 0 | 0 | 0.3 | 0 | 0 | 0 | 1.2 | 0 | 0 | 0 | 14 | 0.2 | 1.2 | 0.1 | 0.1 | 7.9 | 0 | 0 | 0 | 0 | 0 |
| S | 0 | 100 | 3.4 | 0 | 0 | 2 | 0.3 | 0.3 | 0 | 0.6 | 1.1 | 8.9 | 9 | 0.3 | 0.8 | 3.4 | 0 | 0 | 0 | 0 | 0 |
| T | 0 | 0 | 1 | 0 | 0 | 0 | 0 | 0 | 0.2 | 1.2 | 11 | 1 | 0.6 | 3.4 | 0.1 | 0.6 | 7.4 | 0 | 0 | 0 | 0 |
| V | 0 | 0 | 1 | 0 | 0 | 0 | 0.5 | 0 | 100 | 0 | 2.7 | 0 | 0.6 | 1.3 | 0.2 | 0.5 | 1.4 | 0 | 0 | 0 | 0 |
| W | 0 | 0 | 0 | 0 | 0 | 0 | 0 | 0 | 0 | 0 | 0 | 0 | 0.4 | 0 | 0 | 0 | 0 | 0 | 0 | 0 | 0 |
| Y | 0 | 0 | 0 | 0 | 0 | 0 | 0 | 0 | 0 | 0 | 0 | 0 | 14 | 4.2 | 0.1 | 0 | 0 | 0 | 0 | 0 | 0 |

B

| | Sl.No. | Sequence of the loop | % identity at the beginning and ending part | % identity relative to total sequence of the loop | Example organism | Length |
|---|---|---|---|---|---|---|
| ● | | QSPDINQGVDRADPLEQGAGD | 100 | 100 | *E. coli* | 21 |
| ● | 1 | QSPDINQGVDRSDPLEQGAGD | 100 | 95 | *Salmonella enterica* | 21 |
| ● | 2 | QSPDINQGVDRTNPLEQGAGD | 100 | 90 | *Serratia symbiotica* | 21 |
| | 3 | QSPDINQGVEKADPMEQGAGD | 100 | 86 | *Bacteriodota* | 21 |
| | 4 | QSPDINQGIDRTDPREQGAGD | 100 | 86 | *Baumannia cicadellinicola* | 21 |
| | 5 | QSADINQGVERAEPMEQGAGD | 90 | 81 | *Pauldibacter* | 21 |
| | 6 | QSADINQGVDRENPLDQGAGD | 90 | 81 | *Haemophilus influenzae* | 21 |
| | 7 | QSSEIAQGVDRANPLDQGAGD | 80 | 76 | *Caedibacter taeniospiralis* | 21 |
| | 8 | QSPDIAQGVDRDVSNRLEQGAGD | 100 | 74 | *Ectothiorhodospira lacustris* | 23 |
| | 9 | QSADINRGVEREDPMNQGAGD | 90 | 71 | *Bacteroides* | 21 |
| ● | 10 | QSPDINQGVDRAVNDESFEAKANAQGAGD | 100 | 62 | *Chryseobacterium arthrosphaerae* | 29 |
| | 11 | QSPDINQGVERAD--ENADFETKANAQGAGD | 100 | 58 | *Chryseobacterium faecale* | 31 |
| | 12 | QSPDIAQGVDTAHETRVEGKADPLDLQGAGD | 100 | 58 | *Mycolicibacterium* | 31 |
| | 13 | QSPDIAQGVDKALEARQDASDHDPLDLQGAGD | 100 | 56 | *Cellulomonas taurus* | 32 |
| | 14 | QSPDINQGVDRT--LDETDFEARANAQGAGD | 100 | 55 | *Epilithonimonas* | 31 |
| ● | 15 | QSPDIAQGVHLDRNEEDIGAGD | 90 | 55 | *Homo sapiens* | 22 |
| | 16 | QSPDIAQGVDKSIEVRQDSADIDPLDLQGAGD | 100 | 53 | *Cellulomonas* | 32 |
| | 17 | QSPDIAQGVDKSEEERDDASGPSITRDPLDVQGAGD | 100 | 47 | *Luteimicrobium xylanilyticum* | 36 |
| | 18 | QSPDIARGVDTAYERRGGGTAPGGPGDELDRQGAGD | 100 | 44 | *Streptomyces fradiae* | 36 |
| | 19 | QSGDIAQGVN--EALESREGDTDDLSHIGAGD | 80 | 41 | *Streptococcus pyogenes MGAS10394* | 32 |
| | 20 | QSPEIGGTVDRSEEWRQMTPQERARPENTFSEVGAGD | 80 | 35 | *Deinococcus geothermalis DSM 11300* | 37 |

**Figure S6: A**. PSSM of the gatekeeper flexible loop of *Ec*MAT (residue 98-118) using 5000 homologues. The numbers represent the percentage conservation of each residue in each position, that is, 100 indicates a single amino acid is always present at that position in all the sequences, while the number 0 means that the amino acid never appears at that position in any sequence. **B**. 20 different sequences, of the loop of MAT, obtained from BLAST search of *Ec*MAT against 5000 non-redundant sequences. The red letters indicate the mismatched residues with reference to *Ec*MAT gatekeeper helix sequence. The sequences used for loop swap mutations are indicated by red dots.

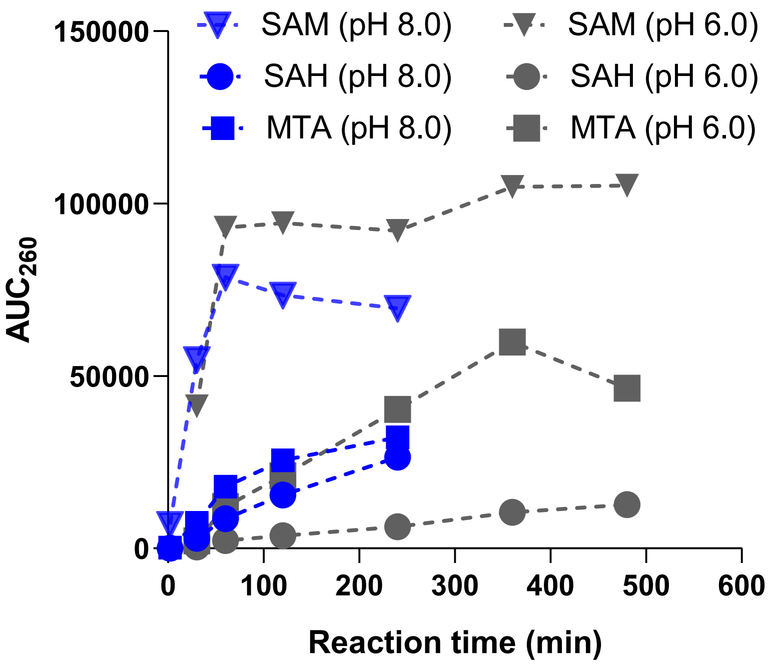


**Figure S7**: A slightly acidic pH decreases the degradation of SAM to SAH, and so pH 6.0 is used for bulk synthesis of SAM and its analogues. The graph shows the $AUC_{260}$ for SAM, MTA and SAH in *Mj*MAT + ATP + methionine reaction mixture injected into HPLC.

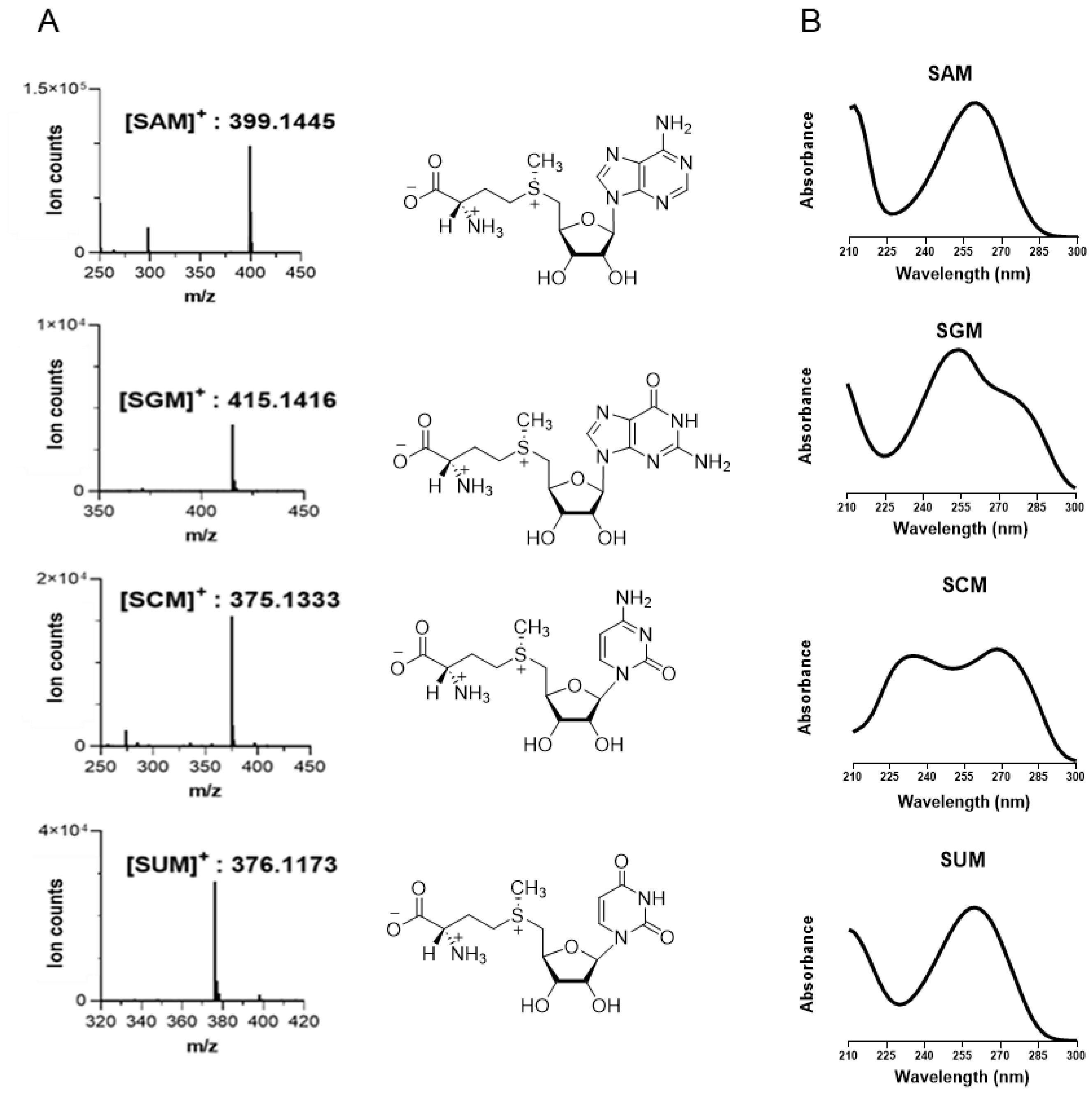


**Figure S8**. The mass fragmentation pattern, structure, and absorbance spectra of SNMs are shown. The expected m/z values are: 399.1445 (SAM), 415.1416 (SGM), 375.1333 (SCM) and 376.1173 (SUM).

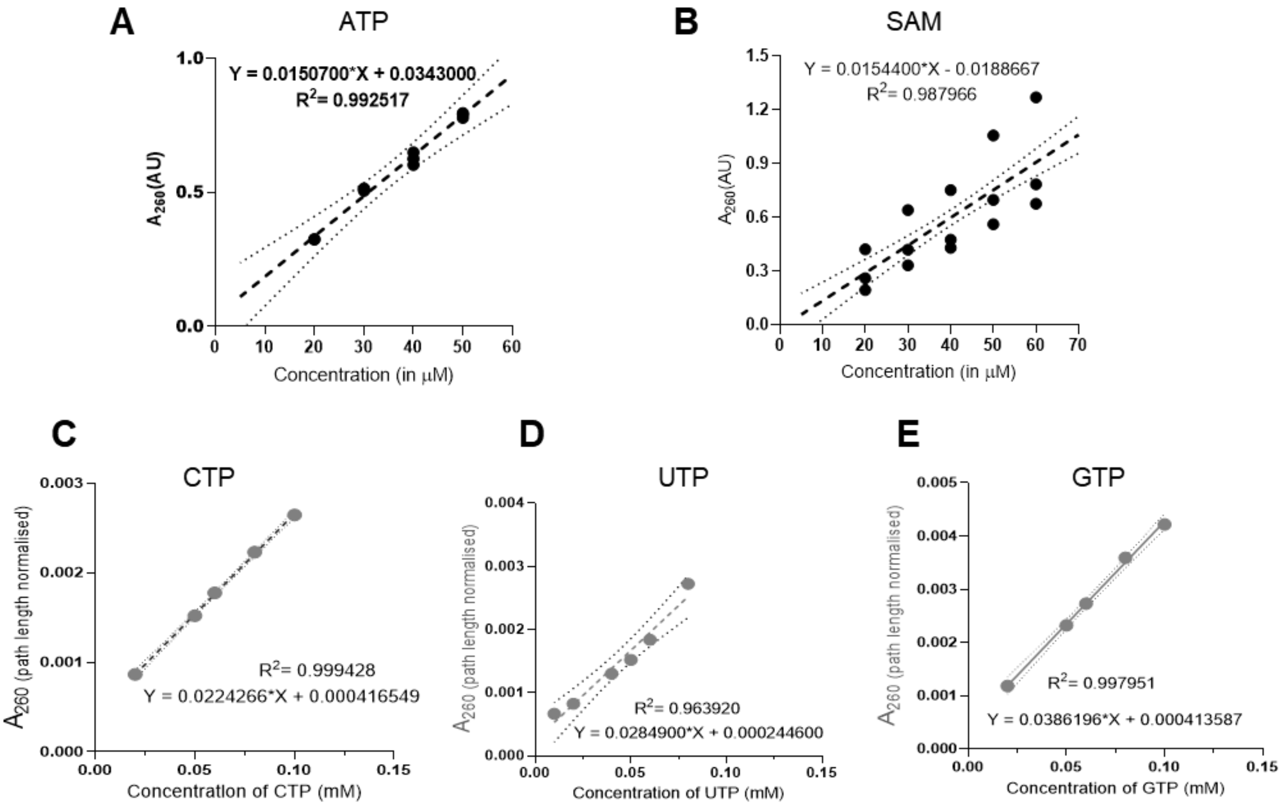


**Figure S9**. Determination of the concentration of SNMs.

Standard absorbance (260 nm) vs concentration calibration curve for ATP (**A**) and for SAM (**B**). The slope gives the molar extinction coefficients (ε) for ATP and SAM as 15070 $M^{-1}cm^{-1}$ and 15440 $M^{-1}cm^{-1}$, respectively. The chromophore being the same in ATP and SAM, and the ε values being similar, we assumed ε values to be equal for other pairs of NTPs and SNMs as well. Standard curves for CTP (**C**), UTP (**D**) and GTP (**E**) are shown.

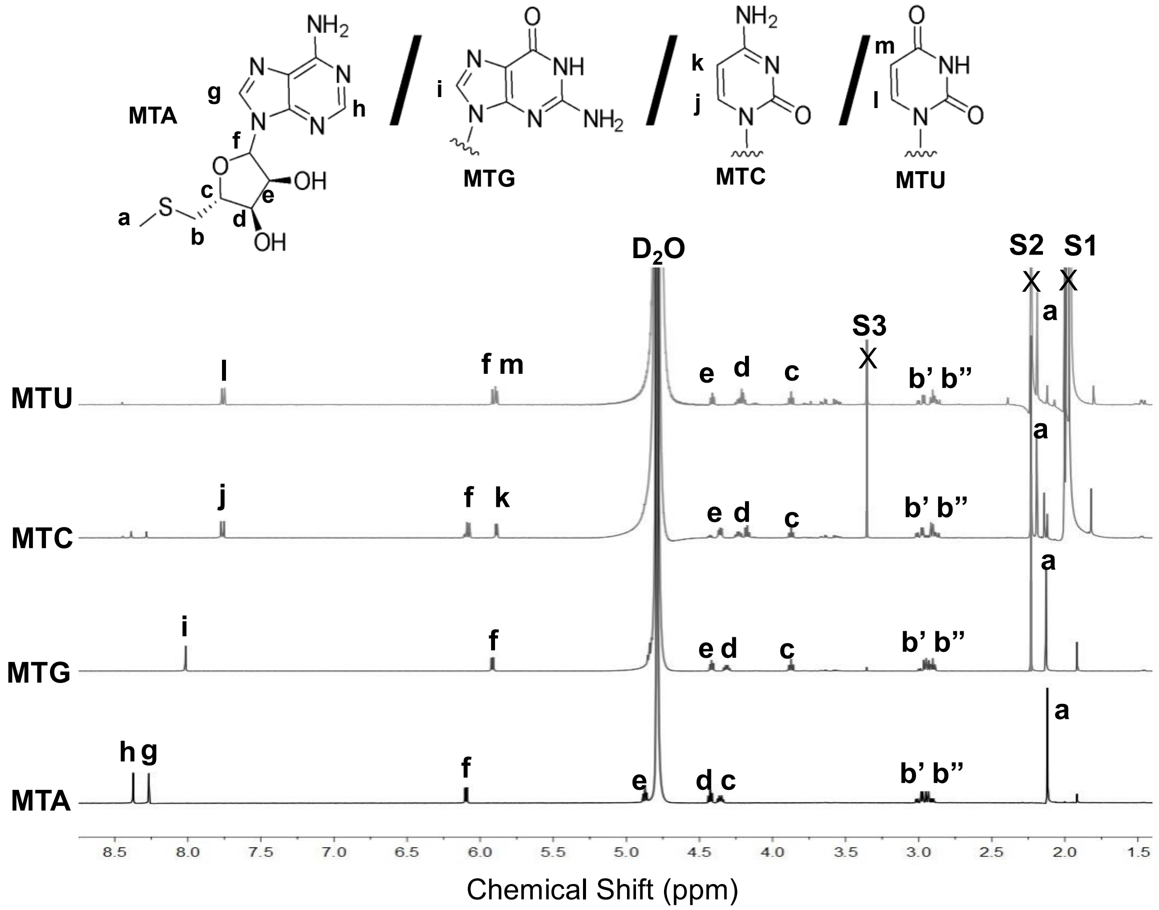


**Figure S10: 1H-NMR spectra of MTA, MTG, MTC, and MTU in $D_2O$.** All four MTNs show peaks (a-f) that correspond to protons in the 5'-methylthioribosyl moieties but vary in peaks g-m that correspond to protons in the nucleobase. Peaks S1, S2 and S3 correspond to the $CH_3$ of residual acetone, acetate, and methanol solvents. The symbols (',") on protons at the same position represent the different shielding due to adjacent chiral carbon. MTNs are collected from HPLC by injecting *Mj*MAT-NTP-methionine reaction mixtures. NMR signals were acquired on a 400 MHz spectrometer.

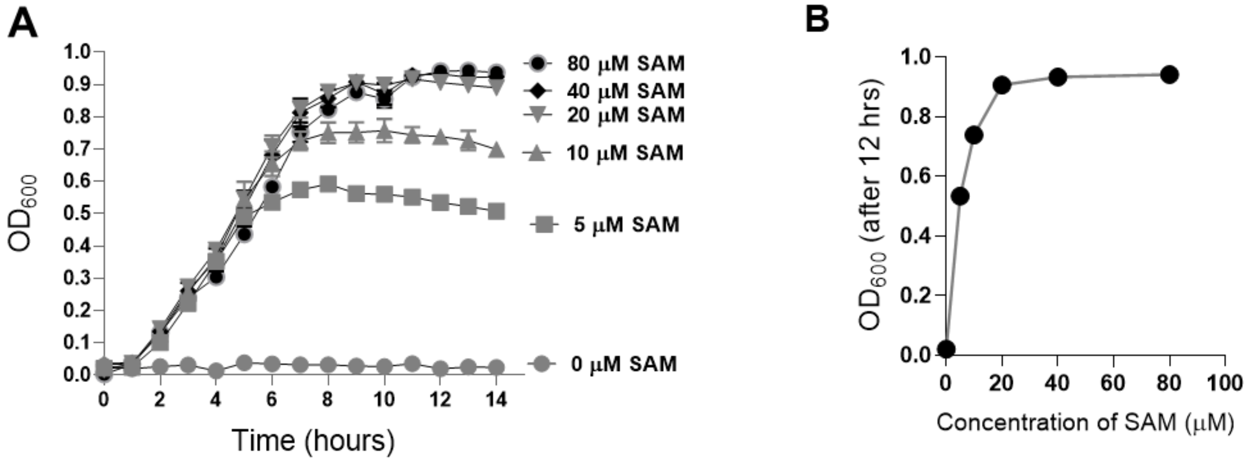


**Figure S11:** MOB1490 growth with SAM titrated in media**.**

**A, B.** MOB1490 growth in LB broth is dependent on the concentration of SAM in the media. Growth saturates beyond 20 μM SAM [5] and so, 50 μM SAM is used for routine MOB1490 growth cultures.

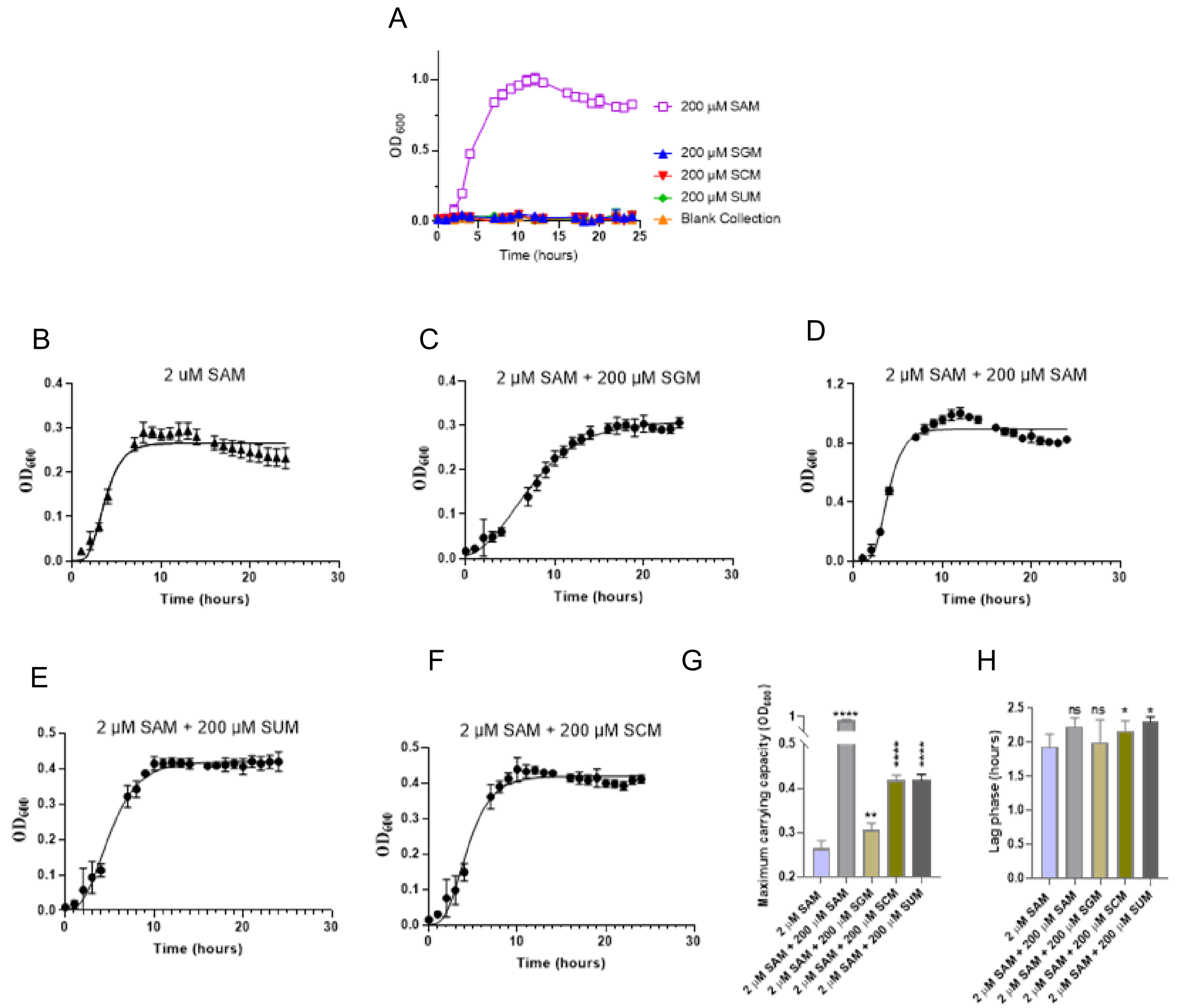


**Figure S12:** MOB1490 growth curve analysis.

**A**. MOB1490 in media, with only SNMs provided, shows no growth. SAM is used as positive control. Blank collection sample is identically processed as that of SNMs except that it is collected from a blank injection in HPLC. **B-H**. The growth curve of MOB1490 in different media supplementations are fitted using Gompertz equation (**B-E**) and three parameters are obtained - maximum carrying capacity (**G**), exponential growth rate and lag phase before exponential phase (**H**). Addition of SGM, SCM or SUM does not significantly alter the lag phase.

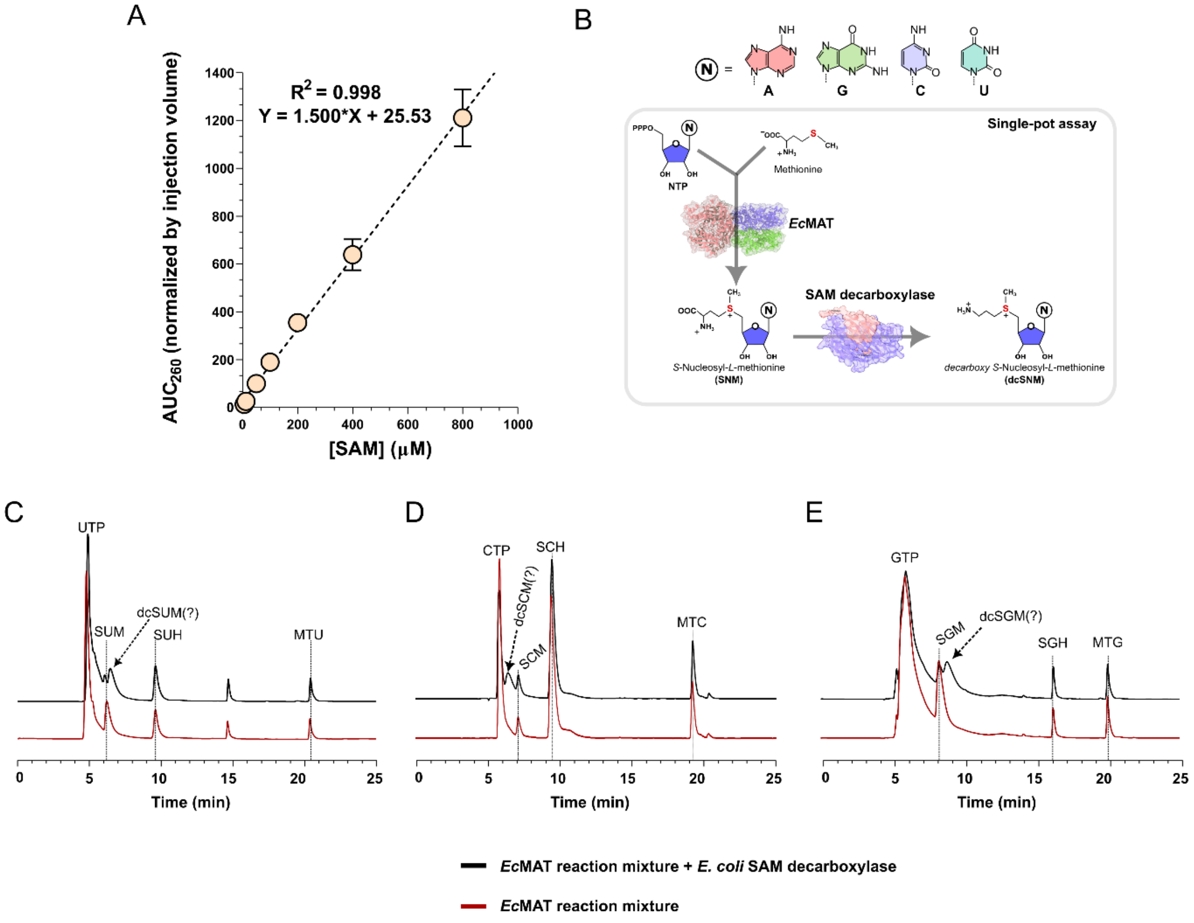


**Figure S13:** Standard curve for SAM quantification using peak area in HPLC chromatograms (**A**). The design of the single-pot two enzyme assay (**B**). HPLC traces of the reaction mixtures of SAM decarboxylase with *in situ* generated SNMs (**C-E**). The reaction mixtures feature an additional peak when SAM decarboxylase is added to the reaction mixtures of *Ec*MAT-N103A with NTPs and methionine. These additional peaks are likely to be for decarboxy SNMs.

| **Sequence (5'-3')** |
|---|
| GTGGATGACTTCTTAAGACAGGAAAACCTGAGTGACGCTTCCCTGGAGGAATGTGAAACCGCCTTAGACCAT CTTATCCAGGGAATACACGGTCTTGACGAACAGGTCATGGCCGCTGTACAAACCCGTTTAGATTCCTTAACCA AACCCCAAGGCAGTTTAGGAGTCCTGGAAGCCCTCGCCAAACAATTGGGCGGGATTCAAGGTCAGCCTTGGC CGGAAGTACAAAAAAAGGCCATTCTCGTCATGGCCGGAGACCACGGCGTGGTGGCGGAAGGGGTCAGTGCC TTCCCTCAGGAAGTAACCCCGCAAATGTTCTATAATTTCCTGGGCGGCGGGGCAGGGATCAATGTCCTGGCC CGTCATGCCGGCGCCGAAGTGATCTGCACCGATGTGGGCATGGCTTTCCCGCTGGATCCGCCCGAGCTTAT GGCTCACCGCATCATGAACGGAACCCATAACATGGCCCAGGGACCGGCTATGAGCCGTAAAGAAGCTCTGCT CTCCCTGCTGACCGGAGCTAAAATCGCCCGCCAGGCTATCGATTCCGGAGTGAATGTCCTGGCCACAGGGG AAGTGGGCATCGGCAATACCACCCCCAGTACAGCCATTATTTCTCTCTTGACAGGTTTGCCGCCCCAGGAGG TTACCGGAAGGGGCACCGGTCTGGATGATGAAGGACTCATCCGCAAGCAAAACGTCATCGCCCGTTCTATCG CCATCAATCAGCCCGATCCTAAGGATGGAGTGGATATCCTCAGCAAAATAGGGGGCTTGGAGATCGGTGCCC TGGCCGGGGCAATTCTGACCGCAGCCTATTCCCGTGTCCCCATCCTTTTAGATGGGGTCATCTCCACTGCCG |

CTGCTCTCATCGCTGTCCGCATCTGCCCTACGGCCCGCTTCTTCCTGATTCCCTCCCATTCTTCTGCGGAGAT
CGGCCATCGAGCCGCCCTGGAACAACTGGAGCTAAAACCCATCATGCATCTGGATTTCCGCTTGGGAGAGG
GAACCGGAGCCGCTATCGCTTTCCATCTTCTGGATGCCTCCATCCGCATTCTCAACGAGATGGCTACTTTTGA
ATCCGCAGGAGTCAGCGGTAAAAACTAAG

**Figure S14:** Sequence of the substrate DNA for *E. coli* DNA methyltransferase assay with the potential methylation sites underlined

### Supplementary material - methods

**Structural alignment of MATs:** To do a structural analysis of MAT, all the crystal structures of MAT available in RCSB PDB were curated using the EC number of MAT (EC 2.5.1.6). We got 120 apo- and holo-MAT structures from 21 different organisms (13 bacteria, 4 archaea and 4 eukaryota). *Ec*MAT (PDB: 7LOO) was taken as a reference. A single sub-unit of the 21 MAT homologues (highest resolution structure of each organism selected) were aligned using the *align* command in PyMOL and the corresponding RMSD values were determined with respect to *Ec*MAT. L147A/I347A *Mj*MAT (PDB ID: 7P84) bound to o-nitro benzyl SAM (ONB-SAM) and *Hs*MAT2a bound to SAM (PDB ID: 8P4H), are considered as two examples of nucleobase promiscuous MAT. The active sites are compared by aligning the PDB structures of *Ec*MAT (PDB: 7LOO), *Mj*MAT (PDB: 7P84) and *Hs*MAT2a (PDB: 8P4H) by aligning the ligands (SAM or ONB-SAM).

**Purification of MAT, SAM Decarboxylase and DAM methylase:** First, the pET15b plasmid carrying the WT or mutant *EcmetK* gene was incorporated into chemically competent *E. coli*

BL21 (DE3) cells via heat-shock transformation. The transformed cells were plated and grown overnight at 37 °C on LB agar plates containing 100 µg/mL ampicillin. Similarly, for overexpression of SAM decarboxylase and DAM methylase, *E. coli* K-12 AG1 strain (obtained from ASKA library) harboring pCA24N-*speD* or pCA24N-*dam (*respectively*)* were grown overnight at 37 °C on LB agar plates having 25 µg/mL chloramphenicol [4].

A single colony was then inoculated into a primary culture containing LB broth and 100 µg/mL ampicillin (for cells expressing MAT) or 25 µg/mL chloramphenicol (for cells expressing SAM decarboxylase or DAM methylase) and grown overnight at 37 °C with 180 rpm shaking. Next day, 1 L LB secondary culture was inoculated with 1% of the primary culture and grown at 37 °C with 180 rpm shaking till the $OD_{600}$ reaches 0.6-0.8. After that, 200 µM (for *Ec*MAT) or 1 mM (for SAM decarboxylase and DAM methylase) isopropyl β-D-1-thiogalactopyranoside (IPTG) was added, and the culture was incubated at 18 °C with 180 rpm shaking for 18 hours, followed by centrifugation at 5500 rpm for 15-30 mins at 4 °C to harvest cells. The harvested cell pellets were stored at -80 °C.

For protein purification, the method was adapted from literature [3,6]. Firstly, the cell pellet was resuspended in lysis buffer (50 mM Tris-Cl pH 8.0, 300 mM NaCl, 0.5 mM phenylmethyl sulfonyl fluoride (PMSF), and 0.025% β-mercaptoethanol (β-ME)). The cell suspension was subjected to lysis by sonication (60% amplitude, 1s on 3s off cycle, 10 minutes). The cell lysate was centrifuged at 18000 rpm for 20 mins at 4 °C, and the supernatant was loaded onto a Ni-IDA/ Ni-NTA column equilibrated with lysis buffer. The column was washed with 10 column volumes of wash buffer (50 mM Tris-Cl pH 8.0, 30 mM imidazole, 300 mM NaCl, 0.025% β-ME). The protein is eluted in elution buffer (50 mM Tris-Cl pH 8.0, 250 mM imidazole, 300 mM NaCl, and 0.025% β-ME). The presence of *the* desired protein in the eluted fractions was verified using SDS-PAGE. Fractions containing the protein were pooled together, loaded onto a Bio-Rad Econo-Pac 10DG desalting column, and then desalted using a desalting buffer (**for *Ec*MAT and SAM decarboxylase**: 50 mM Tris-Cl pH 8.0, 150 mM NaCl, and 0.025% BME; **for DAM methylase:** 20 mM Potassium Phosphate, 200 mM NaCl, 0.025% BME, pH 7.5

balanced using phosphoric acid ). A 12% glycerol solution of the protein was prepared, flash-frozen with liquid $N_2$, and finally stored at -80 °C.

Protein concentration was determined using UV-vis spectrophotometry by measuring the absorbance at 280 nm. The molar extinction coefficients ($\varepsilon$ = 37880 $M^{-1}$ $cm^{-1}$ for MAT, $\varepsilon$ = 39880 $M^{-1}$ $cm^{-1}$ for DAM methylase and $\varepsilon$ = 26360 $M^{-1}$ $cm^{-1}$ for SAM decarboxylase) were predicted using the Expasy Prot Param web tool.
